# Human neurodevelopmental genes housed in massive, ancient gene deserts

**DOI:** 10.64898/2026.03.27.714728

**Authors:** Margaret A. Chapman, Matthew L. Holding, Eirene Markenscoff-Papadimitriou, E. Josephine Clowney

**Affiliations:** Neurosciences Graduate Program, University of Michigan Medical School, Ann Arbor, Michigan, United States of America; Life Sciences Institute, University of Michigan, Ann Arbor, Michigan, United States of America; College of Agriculture and Life Sciences, Department of Molecular Biology and Genetics, Cornell University, Ithaca, New York, United States of America; Department of Molecular, Cellular, and Developmental Biology, University of Michigan, Ann Arbor, Michigan, United States of America; Michigan Neuroscience Institute, University of Michigan, Ann Arbor, Michigan, United States of America

## Abstract

Mammalian genomes contain a remarkable proportion of non-coding sequence relative to protein-coding sequence, including megabase-scale gene deserts. The origin of gene deserts and their function have long been mysteries, and most studies probing these questions have focused on specific deserts linked to transcription factors, like SOX9 and MYC. Here, we describe 21 human “lonely genes” located within some of the largest deserts in the genome, making them exceptionally isolated from other genes in linear sequence space. Together, these regions comprise roughly 3.4% of the human genome. Despite minimal sequence conservation, we find that the sizes and locations of these gene deserts are conserved, and they likely originated at the root of the vertebrate tree. In contrast to the deserts near transcription factors that have been the primary focus to date, we find that most genes housed within these massive deserts, as well as genes on the edges of deserts, are cell adhesion molecules. Additionally, many of the genes associated with these deserts are dysregulated in datasets involving chromatin modifiers linked to neurodevelopmental disorders. Using DNA FISH in mouse olfactory epithelium, we find that lonely genes appose the nuclear lamina. We propose that these ancient regions may play a structural role in the nucleus, and that this structure makes the genes within them uniquely difficult to express. While specific chromatin modifiers are able to access lonely genes, this dependence creates a regulatory vulnerability in brain development.

## Introduction

Genes in mammalian genomes are heterogeneously spaced, with most genes located within a few kilobases (Kbs) of others (Lander et al. 2001; Lindblad-Toh et al. 2005; Venter et al. 2001). This uneven chromosomal distribution is partially attributed to the presence of gene deserts, which are large regions of non-protein-coding DNA which are distinct from telomeres, centromeres, pericentromeres, and subtelomeres (Gregory 2005). Early work on the human genome suggested that roughly ¼ of the human genome could be considered a gene desert, and gene deserts were categorized as either 1) stable deserts which tend to be near genes encoding transcription factors and developmental genes, or 2) variable deserts which tend to be near genes encoding receptors, neurophysiological processes, and intercellular communication (Ovcharenko et al. 2005; Venter et al. 2001). These types of genomic regions are AT-rich relative to the rest of the genome, vary in their degree of conservation, and exhibit a repetitive element profile which is LINE-rich, LTR-rich, and SINE-poor (Ovcharenko et al. 2005).

Despite their prevalence in the genome, the functional role of gene deserts is poorly understood. Consistent with varying conservation, modifications to gene deserts are variably consequential. On one end, several groups have established a link between gene deserts and important regulatory elements, like long range enhancers, which tend to be located around developmental transcription factors (Nobrega et al. 2003; Ovcharenko et al. 2005; Touceda-Suárez et al. 2020). The gene desert around SOX9, for example, has been shown to contain enhancers which are essential for normal craniofacial development (Bagheri-Fam et al. 2006; Gordon et al. 2009; Long et al. 2020). Conversely, deletion of two large deserts in the mouse genome resulted in modest transcriptional changes and didn’t affect animal survival (Nóbrega et al. 2004). Additionally, a gene desert near a well-known oncogene, MYC, has been associated with multiple cancers, and deletion of this desert’s ortholog in mouse reduces breast cancer tumorigenesis (Homer-Bouthiette et al. 2018; Huppi et al. 2012). Together, these data suggest that at best, gene deserts can be oases of critical non-coding, regulatory information, but at worst, they can confer increased risk for disease.

If the impact of these regions is so variable, could there be more than one reason that gene deserts are maintained? Vertebrates have larger genomes than much of the rest of the animal kingdom, despite not gaining a proportionate amount of genic sequence (Gregory 2005). This observation is central to the long-studied C-value enigma, which notes that genome size is highly variable and unrelated to organismal complexity (Gregory 2001). To explain the maintenance of gene deserts, two models have emerged which are not necessarily mutually exclusive. In the first, gene deserts house critical regulatory information in *cis* with the genes they regulate, an arrangement which upholds the structure of the desert (Ovcharenko et al. 2005; Touceda-Suárez et al. 2020). In the second model, gene deserts themselves fulfill a structural role which may be independent of that of linked genes (van Steensel and Belmont 2017). This model is supported by work showing that many gene deserts that are less conserved form lamina-associated domains, or LADs, which are large genomic regions which interact with the nuclear membrane (van Steensel and Belmont 2017). Deletion of core laminar proteins, such as Lamin A and Lamin B, can lead to extensive alterations in gene regulation, and thus, can cause a myriad of diseases (Chang et al. 2022; Harr et al. 2015; Malhas et al. 2007; Poleshko et al. 2013; Shah et al. 2021; Solovei et al. 2013; Worman and Bonne 2007; Zheng and Xie 2019). Most work to date has focused on following individual genes or highly conserved gene deserts, obscuring larger patterns.

It is still unclear how nuclear and genomic organization interact to confer disease risk. It is also unclear how these features have been established over evolutionary time, especially structural features which exist outside the lens of sequence conservation. Additionally, there is little information on the gene deserts that surround genes that aren’t transcription factors. While browsing the human genome, we noticed a set of extremely large gene deserts which contain a single gene, which we call “lonely” genes. In the linear genome, these genes are at least 1 megabase (Mb) away from any other protein-coding gene, and unlike well-studied genes flanking gene deserts (Abassah-Oppong et al. 2024; Bagheri-Fam et al. 2006; de la Calle-Mustienes et al. 2005; Huppi et al. 2012; Tena et al. 2011), ¾ of lonely genes are cell adhesion molecules. We find that over half the genes in this group are enriched for rare variants in individuals with Autism Spectrum Disorder (ASD), and that many show altered gene expression in published datasets of knockout models of chromatin regulators involved in neurodevelopmental disorders (NDDs), such as POGZ and MeCP2 (Boxer et al. 2020; Markenscoff-Papadimitriou et al. 2021; Stessman et al. 2016; Wan et al. 1999; Ye et al. 2015).

To understand whether this unique organization is conserved, making it more likely to be a functional feature rather than an extreme side-effect of a growing genome, we performed comparative microsynteny analysis across diverse vertebrate species’ genomes. We find that these regions are remarkably well-maintained in size and location throughout vertebrates, despite exhibiting minimal sequence conservation. Furthermore, loss or reduction of the lonely organization is most common in lineages that experienced additional whole-genome duplication events and broader genome restructuring. Using DNA FISH in adult mouse olfactory epithelium, a tissue with well-defined nuclear architecture, we find that lonely genes tend to be located near the nuclear periphery. Together, these data lead us to a testable model wherein lonely genes are locked, silent, and at the nuclear periphery unless they are activated by special transcription factors that can penetrate the unique genomic landscape of gene deserts. In sum, we favor the hypothesis that these regions play a bulk structural role in the nucleus, and that the lonely gene organization produces a vulnerability in brain development and tumorigenesis for the genes located inside. Our model leaves unresolved why genes of this type came to be located within such structures.

## Results

### Identifying extremely isolated, “lonely”, genes in the human genome

To formally quantify the heterogeneous distribution of protein-coding genes in mammalian genomes, we collected a list of high-quality, protein-coding gene calls; calculated the distance from each gene to its nearest neighbor (Fig 1A-C); and then performed k-means clustering analyses on these distances. This generated four “distance to nearest neighbor” groups (D, Table 1). The vast majority of genes (∼18,000) are located less than 0.1 megabases (Mb) from their nearest neighbors. In contrast, 21 exceptionally isolated genes are at least ∼1.5 Mb away from any other protein-coding gene (Table 1). The two categories in between contain ∼1000 genes that range from 0.1 to 1.3 Mb away from their neighbors.

**Figure 1:**
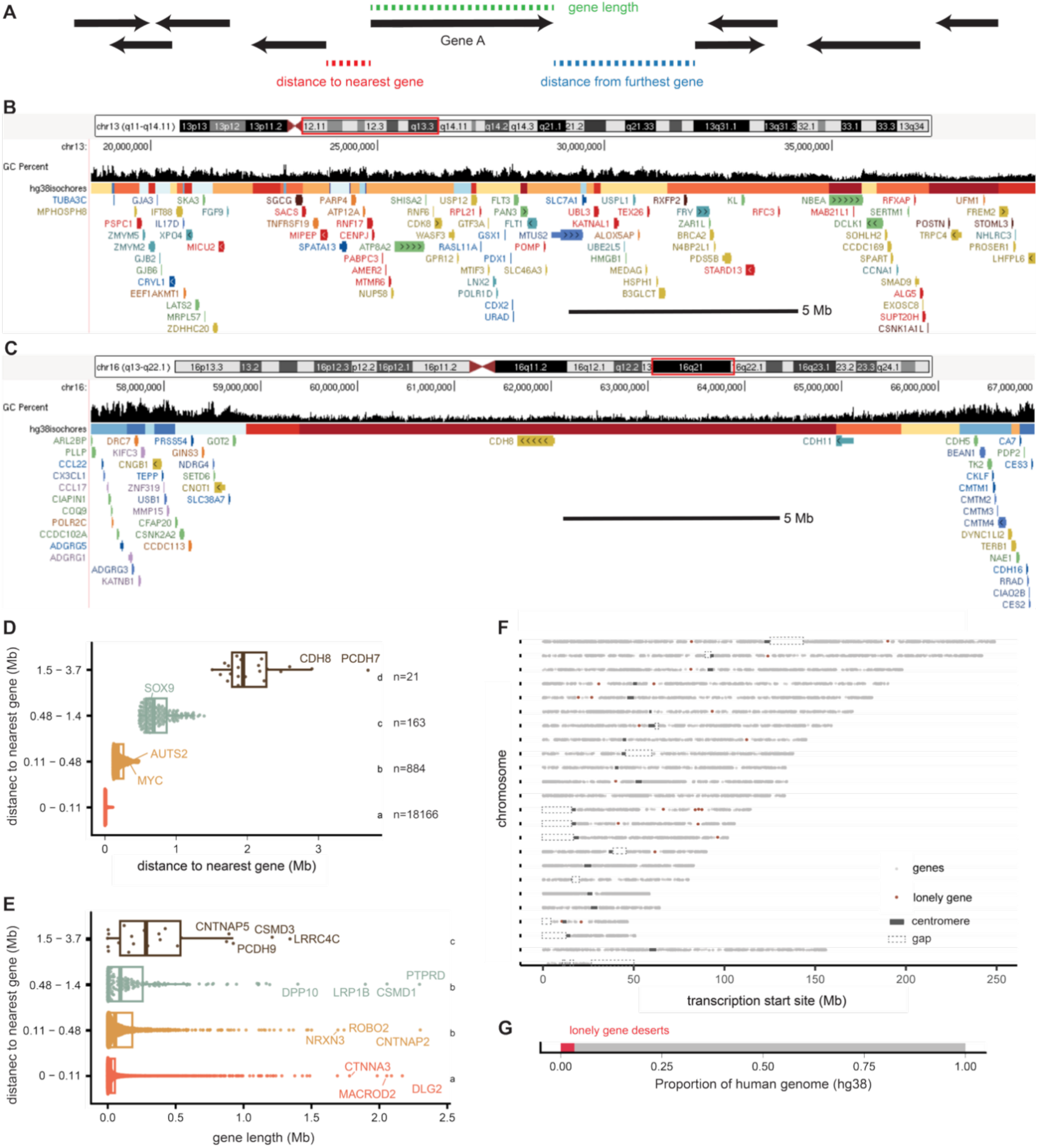
Human lonely genes. A) Genes are indicated by arrows, and the distance to nearest and distance from furthest gene calculations are shown for Gene A. B-C) Screenshots from UCSC Genome Browser display gene density in a typical region of the genome (B) and in a lonely gene desert (C). Isochore GC content is displayed in deciles, with dark red being most AT-rich, yellow being in the middle, and dark blue being the most GC-rich; gene model colors reflect the AT content characteristics of the gene itself (Brovkina et al. 2023). D) Distance to nearest gene calculations are grouped by k-means clustering, with the most isolated, “lonely” genes shown in brown. The number of genes in each group is labeled for each cluster as “n =”. E) Gene length distributions for gene loneliness categories as in (D). Letters indicate significance from a pairwise Wilcoxon test with FDR correction. Significantly different groups carry different letters. F) Every protein-coding gene in the human genome is displayed by the location of its TSS, where each row is a chromosome, centromere sequence is noted by grey blocks, and gaps are notated by lighter grey, dashed boxes. Lonely genes are colored red. G) Proportion of the human genome that is consumed by the 21 lonely gene deserts.

**Table 1:** Lonely gene description. Each row shows characteristics associated with each lonely gene. Distance to nearest neighbor calculation is described above and presented here in Mb, colored by loneliness relative to other genes, with most lonely genes darker green. Gene families were pulled from UniProt. Whether or not a gene is associated with cell adhesion was determined by a manual literature search. SFARI gene classifications for autism spectrum disorder association (ASD) are as follows: category 1 = high confidence, category 2 = strong candidate, category 3 = suggestive evidence. Full descriptions of each category are available at SFARI GENE’s online About page (Abrahams et al. 2013). POGZ and MeCP2 regulation from Markenscoff-Papadimitriou *et al*. 2021 and Tenreiro *et al*. 2025 as well as gene length clusters were determined as described below (Markenscoff-Papadimitriou et al. 2021; Tenreiro et al. 2025).

| Gene name | Chrom | Distance to nearest neighbor (Mb) | Gene family | Cell adhesion? | SFARI categorization | Effect of POGZ loss | Effect of MeCP2 loss | Gene length (Kb) |
| --- | --- | --- | --- | --- | --- | --- | --- | --- |
| ADGRL2 | chr1 | 1.87 | adhesion G protein-coupled receptors | Yes |  |  |  | 193 |
| BRINP3 | chr1 | 1.68 | BMP/retinoic acid inducible neural specific |  | category 3 |  | Down | 380 |
| LRRC4C | chr11 | 1.85 | Ig-like cell adhesion molecule family | Yes | category 1 | Down |  | 1345 |
| PCDH9 | chr13 | 2.47 | non-clustered protocadherin | Yes | category 2 |  | Down | 928 |
| SLITRK1 | chr13 | 1.91 | SLIT and NTRK like family | Yes |  | Down |  | 5 |
| SLITRK5 | chr13 | 1.87 | SLIT and NTRK like family | Yes | category 2 | Down |  | 8 |
| SLITRK6 | chr13 | 1.87 | SLIT and NTRK like family | Yes |  |  |  | 7 |
| FLRT2 | chr14 | 2.23 | fibronectin type III domain containing | Yes |  | Down | Down | 124 |
| LRFN5 | chr14 | 2.17 | fibronectin type III domain containing, Ig-like cell adhesion molecule family | Yes | category 1 | Down | Down | 298 |
| NR2F2 | chr15 | 1.62 | nuclear hormone receptor |  |  |  |  | 10 |
| CDH8 | chr16 | 2.91 | type II classical cadherin | Yes | category 2 | Down | Down | 389 |
| CNTNAP5 | chr2 | 1.73 | contactin associated protein family | Yes | category 2 |  |  | 896 |
| NCAM2 | chr21 | 2.59 | fibronectin type III domain containing, Ig-like cell adhesion molecule family | Yes |  | Down | Down | 545 |
| GBE1 | chr3 | 1.72 | glycosyl hydrolase |  | category 2 |  |  | 272 |
| ADGRL3 | chr4 | 2.2 | adhesion G protein-coupled receptors | Yes |  |  | Down | 878 |
| PCDH7 | chr4 | 3.69 | non-clustered protocadherin | Yes |  | Down |  | 426 |
| CDH18 | chr5 | 1.76 | type II classical cadherin | Yes |  |  | Down | 517 |
| CDH9 | chr5 | 2.24 | type II classical cadherin | Yes | category 2 |  | Down | 158 |
| EPHA7 | chr6 | 2.16 | EPH receptors | Yes |  | Down |  | 180 |
| POM121L12 | chr7 | 1.51 | POM121 transmembrane nucleoporin like |  |  |  |  | 1 |
| CSMD3 | chr8 | 1.97 | sushi domain containing |  | category 3 |  | Down | 1214 |

**Figure S1:**
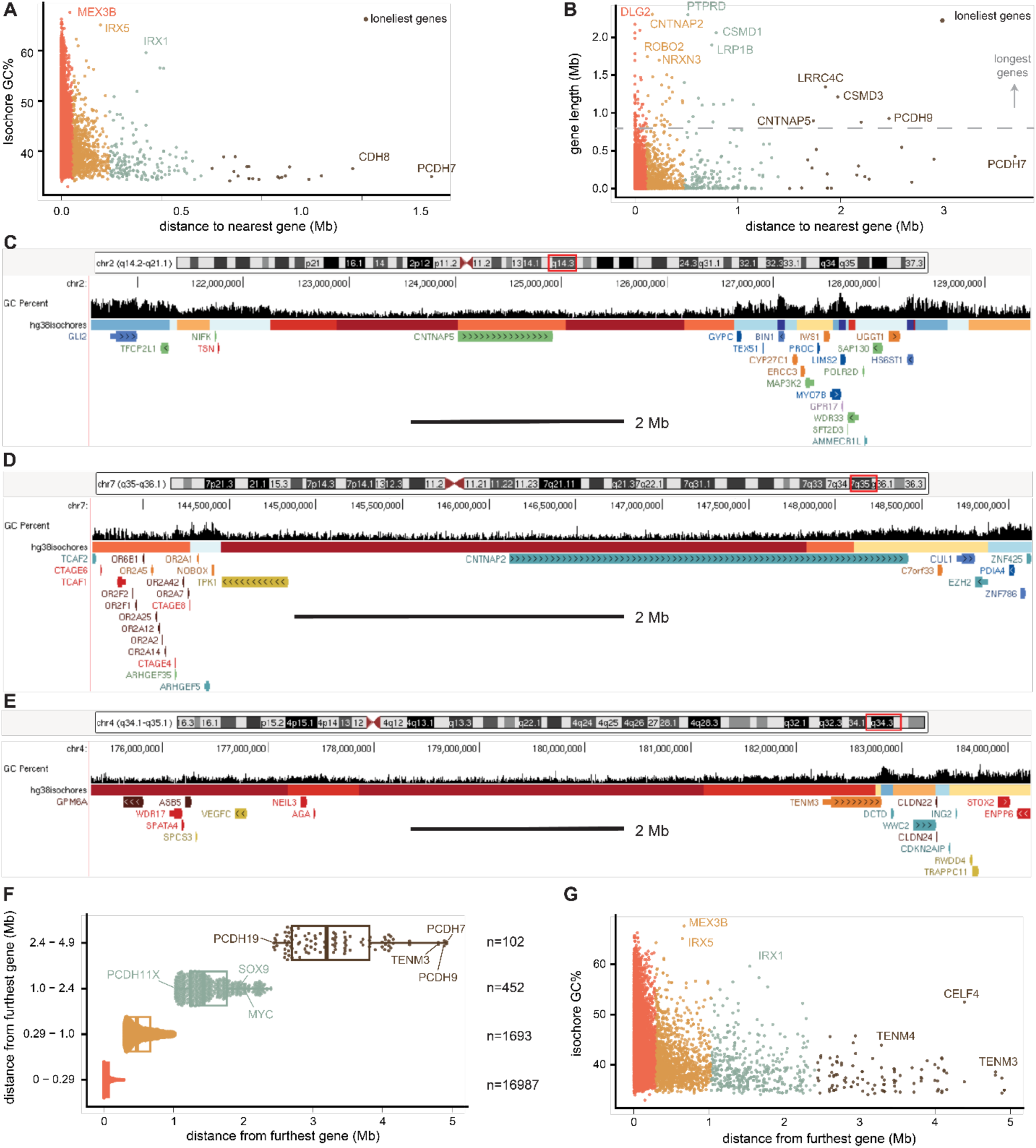
A) The GC content of the isochore containing each gene, as calculated from previous work, is displayed against distance to nearest gene calculations, colored by distance clusters (Brovkina et al. 2023). B) Gene length is displayed against distance to nearest gene, with genes in the longest gene cluster indicated above the dashed line. C-E) The UCSC Genome Browser screenshots display a long and lonely gene (C), a paralogous long gene that is not lonely (D), and a gene desert without a gene (E). F) “Furthest distance to gene” clusters are displayed, with “n=” indicating the number of genes in each category. G) Isochore GC% is displayed against distance from furthest gene calculations, colored by furthest distance clusters.

Consistent with previous reports of gene desert sequence content, the deserts containing the loneliest genes are exceptionally AT-rich (Fig S1A) (Holmquist 1992; Ovcharenko et al. 2005; Versteeg, et al. 2003). We plotted every gene in the human genome at the chromosome level (Figure 1F) and observed that these regions were visually identifiable and distinct from centromeres, pericentromeres, and telomeres. Many studies have focused on “long genes,” i.e. transcribed units of unusual length. We found that the transcribed units of genes located in deserts, measured from TSS to TES, are longer than the average gene, but include a broad range of gene sizes (Fig 1E and S1B). Moreover, many long genes are not lonely (Fig S1D). (These observations are not a side effect of our metric, since we calculated “distance to neighbor” from either side of the gene, not from a midpoint (Fig 1A, S1C-D).) To highlight their unique genomic organization and distinguish them from long genes, we refer to genomically isolated genes as “lonely genes.” By our definition, lonely genes have no neighbors on either side. Measuring each gene’s distance to its *furthest* neighbor, rather than the nearest neighbor, picks out a related set of 100 genes that have a large desert (2.5-5Mb) on at least one side (Fig S1E-F). Similar to the lonely genes, these deserts are also almost entirely in the bottom 10% of genome-wide GC content (Fig S1G).

### Characterizing lonely genes

Out of 21 lonely genes, 16 (∼75%) are predicted to be cell adhesion molecules, and these genes, along with those that have a large desert on one side, but not both, also function in synapse organization and assembly (Fig 2A, Fig S2A, Supplementary Table 1). Thus, these genes may be particularly important for the developing brain. Given the enrichment of neurodevelopmental functions based on gene ontology (GO), we asked if these genes were explicitly associated with increased risk of any neurodevelopmental disorders (NDDs) (Dekker 2007; Li et al. 2020; Wang and Willard 2012). Alleles of 10 of the 21 lonely genes (∼50%) have been linked to autism, compared to 25%, 15%, and 6% in less isolated clusters (Fig 2C, Table 1) (Banerjee-Basu and Packer 2010).

**Figure 2:**
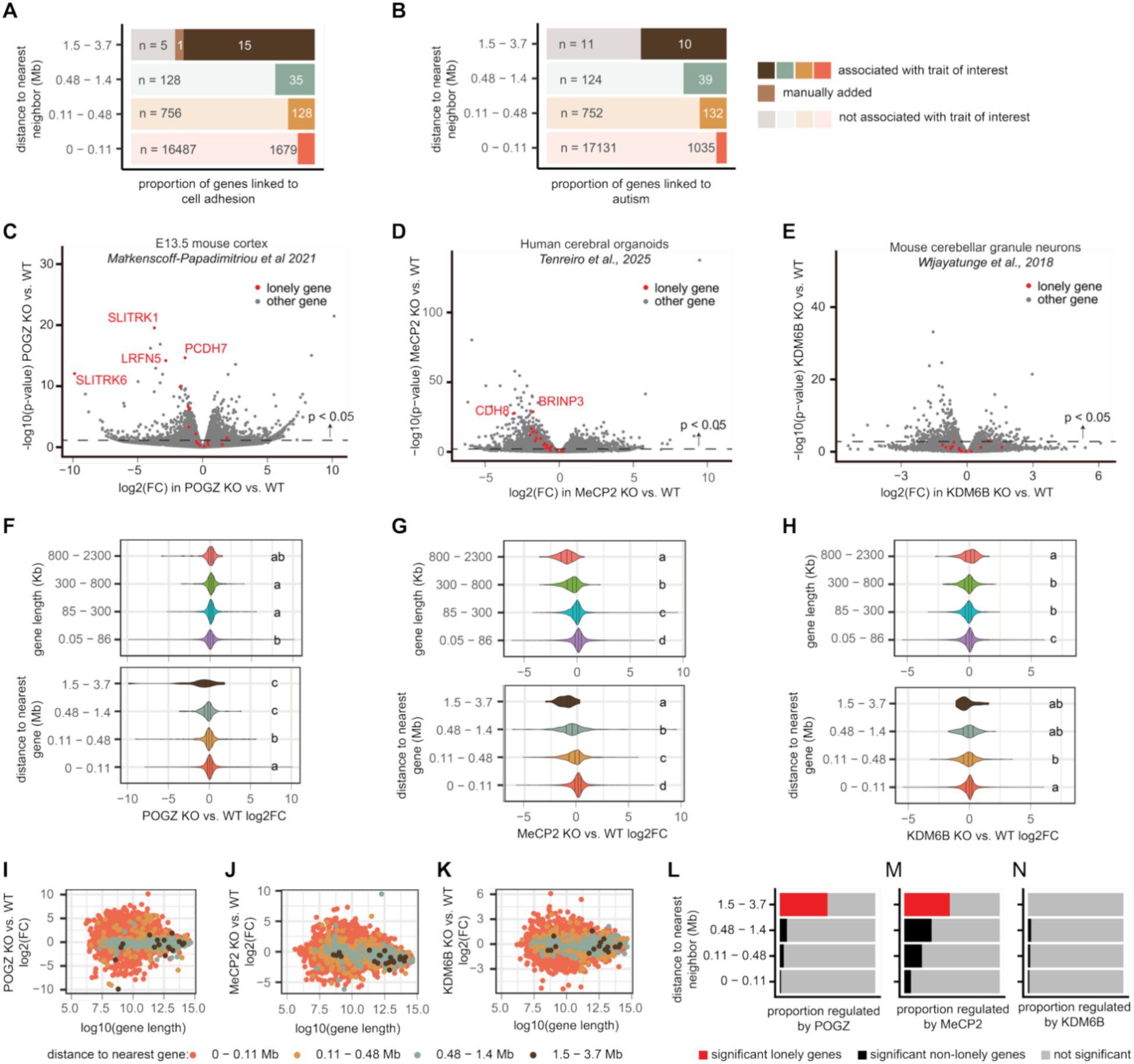
Lonely genes are linked to neural development. A) Each row corresponds to a distance to nearest gene cluster, and darker shading represents the proportion of genes in that group that are linked to cell adhesion. B) Each row indicates the proportion of genes linked and not linked to autism spectrum disorder in the SFARI, grouped by distance to nearest gene. SFRARI gene classifications are as follows: category 1 = high confidence, category 2 = strong candidate, category 3 = suggestive evidence (Abrahams et al. 2013). Full descriptions of each category are available at SFARI GENE’s online webpage. C-E) Re-analysis of gene expression data from neurons lacking chromatin regulators POGZ, MeCP2, and KDM6B (Markenscoff-Papadimitriou et al. 2021; Tenreiro et al. 2025; Wijayatunge et al. 2018).Log2(fold change) and log10(p-value) of knock-out (KO) versus wildtype (WT) gene expression is displayed. Red genes are lonely genes, and genes above the dashed line had p < 0.05. For a gene to be considered significantly regulated, had to have both a p-value < 0.05 and a log2FC > 1. F-H) KO vs. WT log2(FC) by both gene length and distance to nearest gene clusters. Different letters indicate significant differences between groups, as determined by pairwise Wilcoxon test with FDR correction. I-K) Log2(FC) of KO vs. WT compared to log10(gene length) is displayed for each gene, with colors corresponding to distance to nearest gene cluster. L-N) Each row displays the number of significantly regulated genes in each “distance to nearest” gene cluster as determined by two-proportion Z test. Red indicates significantly regulated lonely genes from (C-E). Genes we call as “significantly regulated” have a p-value < 0.05 and a log2FC > 1.

**Figure S2:**
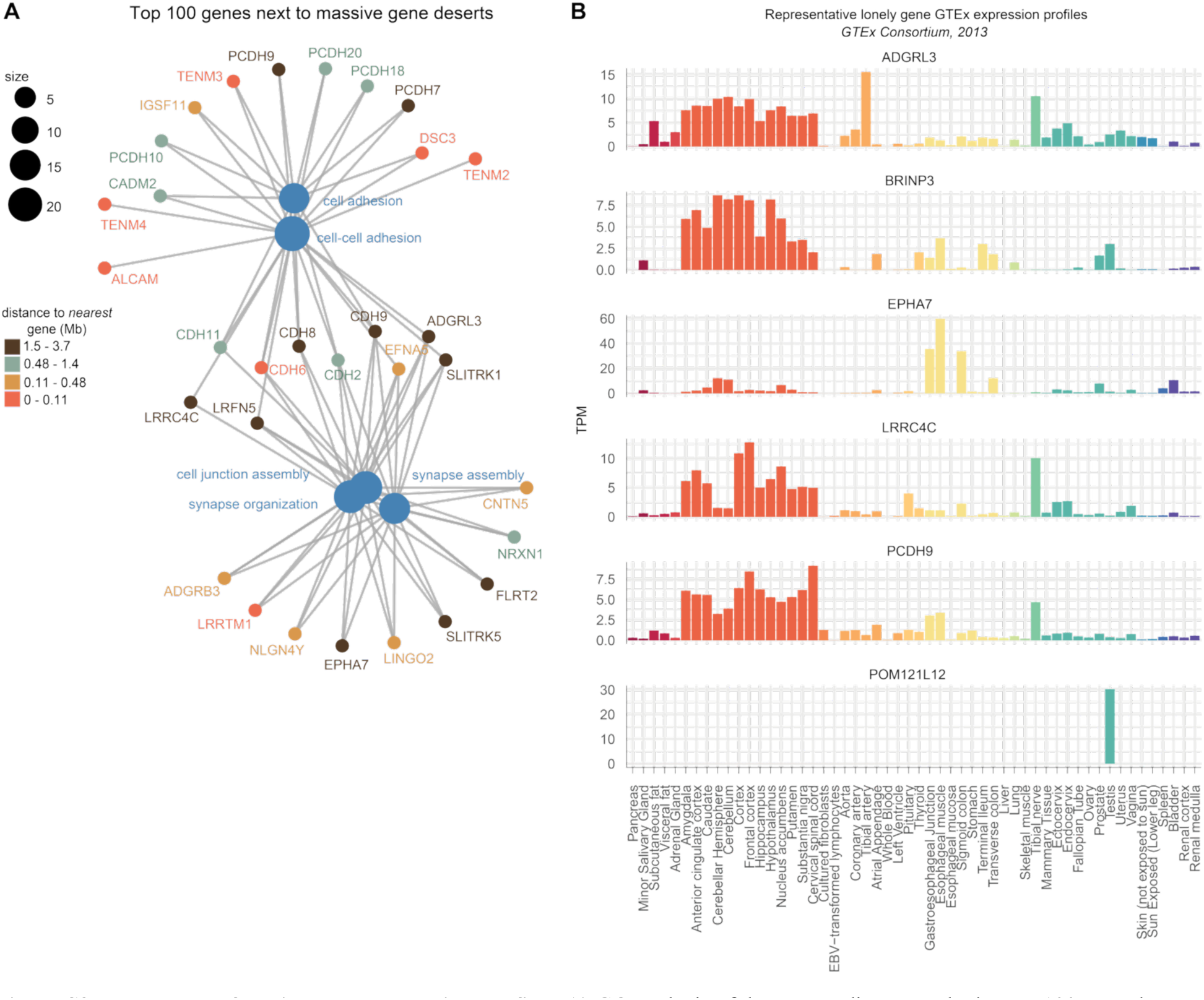
Lonely gene functions and expression profiles. A) GO analysis of the genes adjacent to the largest 100 gene deserts in humans, as pulled from the “distance from furthest” gene calculation. This captures both lonely genes and genes on the edge of a large desert. Gene labels are colored according to which “distance to nearest” group they belong to. B) Bulk tissue-level gene expression for 6 representative lonely genes, from GTEx (GTEx Consortium 2013). Tissues are colored by organ or organ systems (ie: brain regions are displayed in orange, and GI tract elements are displayed in yellow).

Interestingly, only one gene in the most isolated set is neither a cell adhesion molecule, nor associated with autism. Expression data from GTEx shows that this gene, POM121L12, is only expressed in testis, unlike the others, which are variably expressed in other tissues (Supp Fig 2B) (GTEx Consortium 2013). *De novo* genes are typically only transcribed in the testis due to widespread transcription and particularly permissive expression landscape (Begun et al. 2006; Kaessmann 2010; Kondo et al. 2017; Levine et al. 2006; Witt et al. 2019; Xia et al. 2020). Since POM121L12 is also far less conserved than other genes, this suggests POM121L12 may be a new gene (Fig S3H).

Previously, we observed that small genomic clusters of synaptic genes, such as SLITRK1, SLITRK6, SLITRK5 family on human chromosome 13, depend on POGO transposase derived zinc finger TF, POGZ, for their expression (Markenscoff-Papadimitriou et al. 2021). Here, we noticed that many of these synaptic genes that depend on POGZ for their expression are also members of the lonely gene set: POGZ knockout in E13.5 mouse cortex results in downregulation of a significant proportion of lonely genes (8/21): EPHA7, LRRC4C, SLITRK1, SLITRK5, LRFN5, FLRT2, CDH8, and NCAM2 (Table 1, Fig 2C, F, I, and L, χ^2^ = 310.98, df = 3, p < 2.2e-16). Since many lonely genes are also long, we wondered if POGZ was also disproportionately regulating long genes, but we did not find this to be true in this tissue, despite identifying a significant interaction between gene length and loneliness (Fig 2F and I, Supplementary Table 2).

Since heterozygosity of POGZ is causal for the syndromic autism spectrum disorder (ASD) White-Sutton Syndrome, we wondered if other chromatin modifiers linked to NDDs or ASD also play a special role in the regulation of genes with this architecture (Assia Batzir et al. 2020). Using bulk RNA-sequencing data generated by Tenreiro et al, we find that in human cerebral organoids, MeCP2 regulates several lonely genes (Tenreiro et al. 2025)(Fig 2D, G, J, and M, χ^2^ = 266.6, df = 3, p-value < 2.2e-16). However, unlike with POGZ, this trend is dominated by the long genes in the lonely set, a feature which is known to enriched in MeCP2-regulated genes (Fig 2G, J, and M, Supplementary Table 2) (Boxer et al. 2020; Gabel et al. 2015; Sugino et al. 2019; Tenreiro et al. 2025). We also find that KDM6B, another neuronal chromatin regulator, is important for expression of long genes, but not lonely genes (Wijayatunge et al. 2018) (Fig 2E, H, K, and N), χ^2^ = 7.412, df = 3, p-value = 0.060).

### Are human lonely genes always lonely in other species?

If lonely genes are involved in cell adhesion and brain development, does their extreme organization in the human genome protect them or produce a vulnerability in brain development? To understand whether this organization is unique to the human genome (i.e. perhaps more likely to be an anomaly), we asked whether the organization of these genes is conserved. To do this, we tracked microsynteny (gene order) both manually and in Genomicus, following each lonely gene and its associated desert as far back as the lonely gene and/or its neighboring genes could be identified (Fig 3 and Fig S3A-H) (Louis et al. 2015; Louis, Muffato, and Roest Crollius 2013; Nguyen et al. 2018, 2022). In almost all vertebrates, we found enormous gene deserts surrounding orthologs of human lonely genes, except POM121L12, the testis-expressed gene that we hypothesized evolved recently (Fig S3H). In many cases, microsynteny was retained, such that the gene on the other side of the enormous desert was also orthologous to that found in human. In birds and reptiles, lonely gene deserts are smaller, but they are proportionally about as large as mammalian gene deserts relative to the smaller genome sizes among Sauria (Fig 3 and Fig S3F).

**Figure 3:**
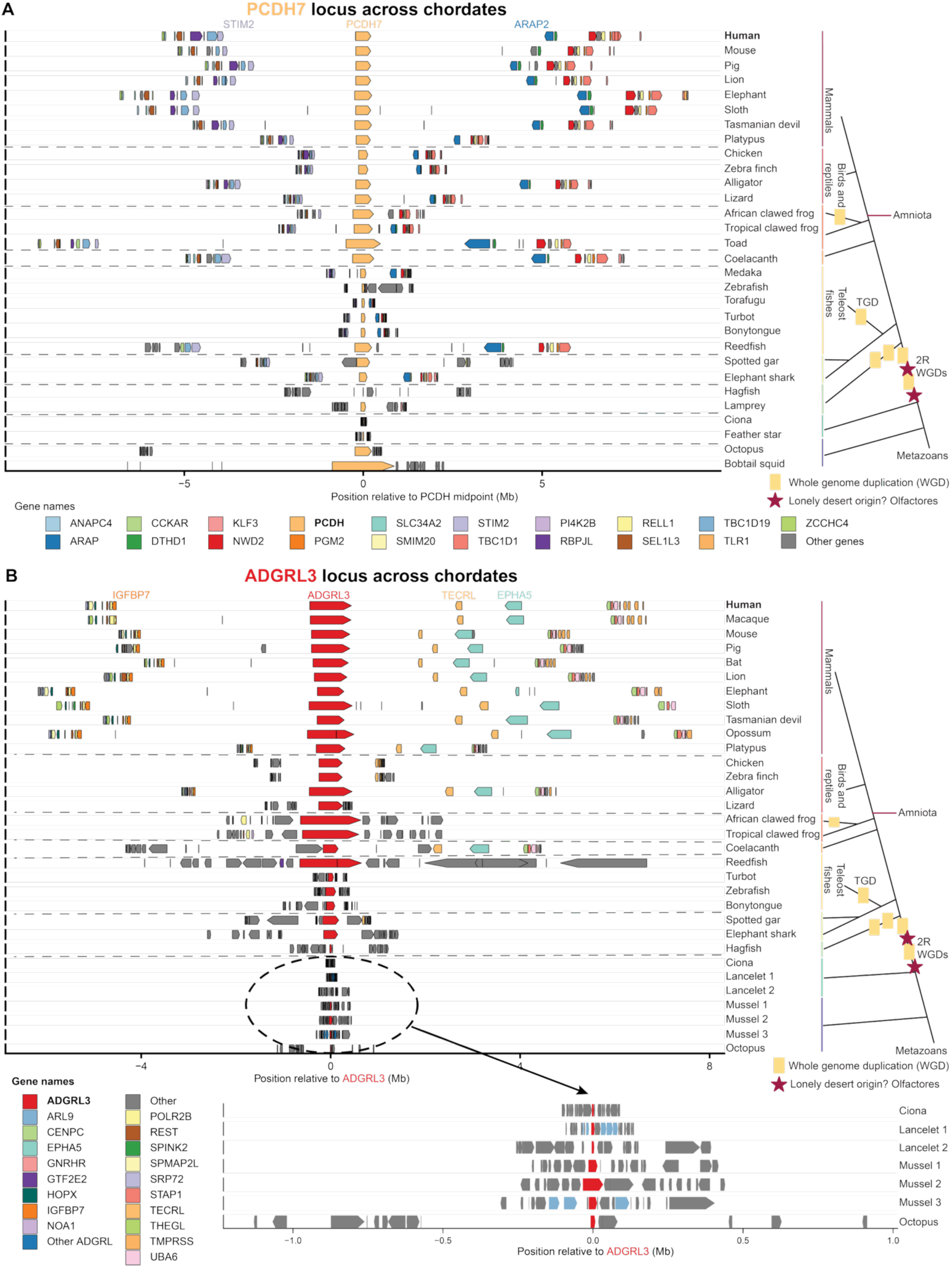
Lonely gene desert size across Metazoans. A) PCDH7, (beige arrow), and surrounding syntenic genes are displayed here. Orthologs of genes that share microsynteny with the locus in human are colored, as indicated. Genes that are not syntenic with the human locus are shown in grey. Yellow blocks on the tree diagrams represent whole genome duplications. B) Similarly, ADGRL3 is displayed in red for each species, and surrounding syntenic genes are also colored. The inset displays ADGRL3 loci in a subset of non-vertebrate genomes. Red stars indicate presumed lonely gene desert origin.

**Figure S3:**
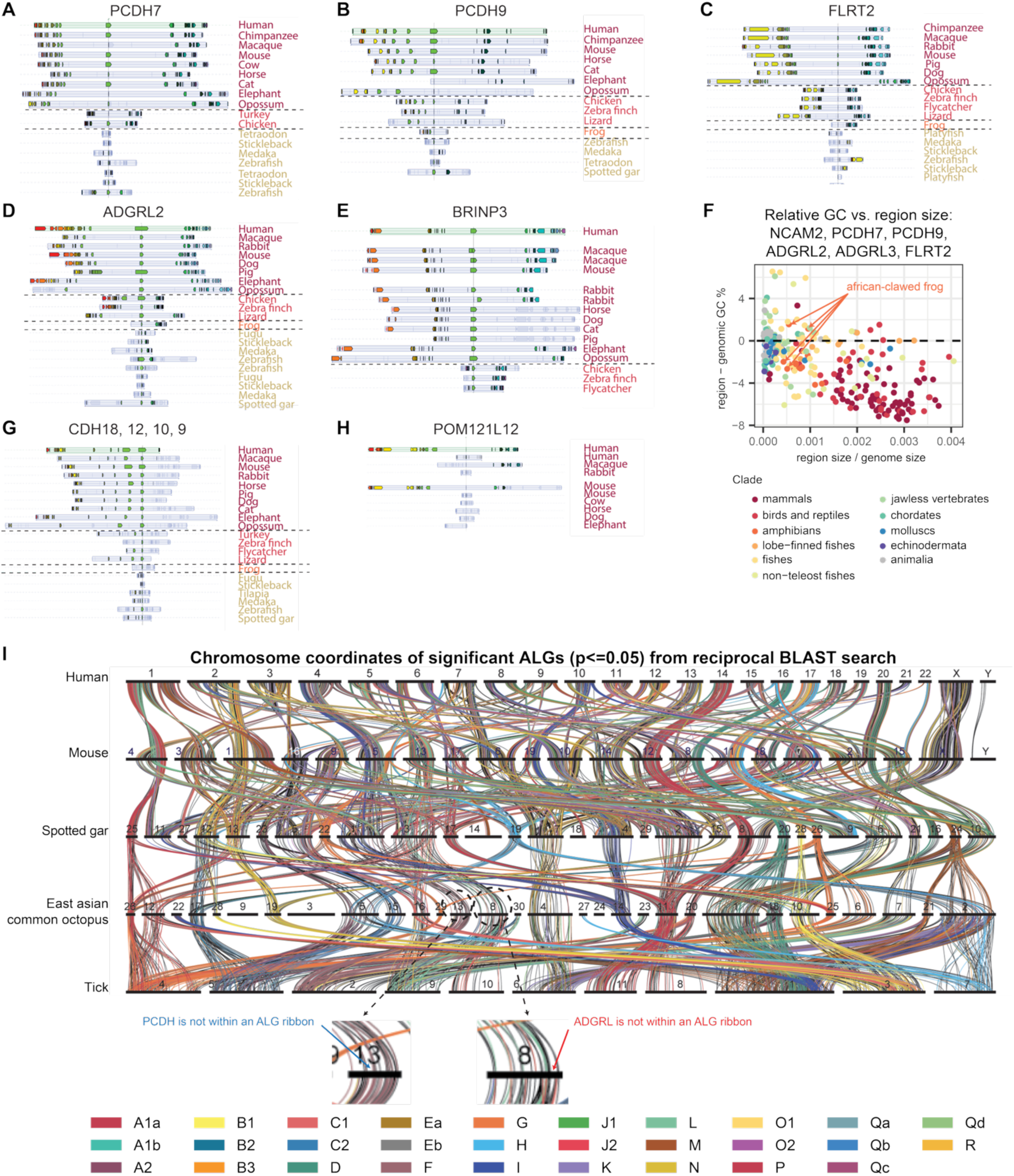
Lonely gene spacing across Metazoans. A-E, G-H) Screenshots from Genomicus are displayed for several representative lonely gene loci. Each row corresponds to one species. Center line and associated green arrow indicate the gene of interest, and other colored genes are indicative of homologous genes across species. Species names are colored based on their respective clades as indicated in F. F) Region GC relative to genomic GC is displayed against locus lengths controlled by genomic GC. For each species, up to six dots are shown, corresponding to the 6 lonely loci as indicated (NCAM2, PCDH7, etc); dots are colored by the species’s clade. I) A ribbon plot, generated by ODP, shows syntenic blocks, or ancestral linkage groups (ALGs), that are occurring more often than would be expected by random shuffling. Each black bar represents a chromosome from one of 5 species: human, mouse, spotted gar, octopus, and tick (Schultz et al. 2023).

There were a few additional species where large gene deserts were observed in lonely gene homologs despite loss of microsynteny and extensive evolutionary distance: East Asian common octopus (O. sinensis), California two-spot octopus (O. bimaculoides), and bobtail squid (E. scolopes) (Fig 3). To track whether or not these deserts were convergent (ie: separately evolved) or conserved, we performed reciprocal BLAST searches and identified conserved lineages between 3 genomes with known synteny (human, mouse, and spotted gar), O. sinensis, and a genome without a lonely protocadherin (tick) using Oxford Dotplot (Fig S3I, Supplementary Table 6) (Schultz et al. 2023). We successfully identify lonely PCDH’s and ADGRL’s when comparing vertebrates with known, conserved lonely gene deserts (ie: mouse, human, and spotted gar; Supplementary Table 6). Reciprocal BLAST identifies the O. sinensis genes displayed in Figure 3 as orthologs to human PCDH9 and ADGRL3, but we do not find them in linkage groups with vertebrates (Supplementary Table 7). This suggests that the organization of these regions arose convergently in the cephalopod and vertebrate clades, rather than having a common origin, but we can’t rule out the possibility that they are conserved and have lost synteny over evolutionary time.

In three species or clades, lonely genes were sometimes not located in gene deserts: African-clawed frog, teleost fishes, and jawless vertebrates (Fig 3 and Fig S3H). Interestingly, the genomes of each of these species have undergone species- or clade-specific whole genome duplications (WGDs) (Furman et al. 2018; Glasauer and Neuhauss 2014; Kobel and Du Pasquier 1986; Marlétaz et al. 2024; Mehta et al. 2013; Nakatani et al. 2021; Putnam et al. 2008; Session et al. 2016; Simakov et al. 2020; Tymowska and Fischberg 1973) (Fig 3, Fig S3, Supplementary Table 3). We note that two rounds of whole genome duplication (2R WGD) are thought to have occurred at the root of the vertebrate tree, so all vertebrates have undergone at least 2 WGDs (Dehal and Boore 2005; Marlétaz et al. 2024; Ohno 1970).

To examine this trend, we sorted genomes into three categories: those without WGDs (ie: a subset of protostomes), those with two (ie: most vertebrates that aren’t teleost fish), and vertebrates with 3 or more. Here, a clear pattern emerges, whereby those that underwent the vertebrate 2R WGD events carry the largest deserts, vertebrates that underwent additional duplication events (a total of 3 or more) in the middle, and those without a known WGD rarely have a discernable gene desert (Fig S4B). Together, these data lead us to the hypothesis that the lonely gene deserts we identified in mammals emerged around the origin of vertebrates, and that they were variably lost in vertebrate species which underwent subsequent clade-specific WGD events. As we describe above, we cannot completely rule out an earlier origin of some of these deserts and their subsequent loss in non-vertebrate, non-cephalopod branches.

Next, we asked if this organization arose before or after the 2R WGDs at the root of vertebrates. We noticed that several genes in the lonely set are paralogs (ie: PCDH7 and 9) and that several are likely to be a special kind of paralog that results from WGD, called ohnologs. Analyses by Sacerdot *et al*. show that the majority of lonely genes fall into two categories: those highly likely to be ohnologs (i.e., members of gene sets that resulted from 2R duplications, Fig. 4A–G) and those with a lower but notable probability of being ohnologs (Fig. 4H–M) (Sacerdot et al. 2018). We therefore asked whether the predicted ohnologs of lonely genes share a similar organization, which would indicate that the organization arose before they were duplicated during the 2R events. A “perfect” ohnolog set contains four genes, on four separate chromosomes, corresponding to a single gene that was present before the 2R duplications. While the 2R whole-genome duplication model remains disputed (Smith et al. 2018; Smith and Keinath 2015), the origin of ohnologs through large-scale duplications is clear. Whether because of duplication events that didn’t affect the whole genome or due to subsequent gene gains and losses, many ohnolog sets do not have exactly four members.

**Figure 4:**
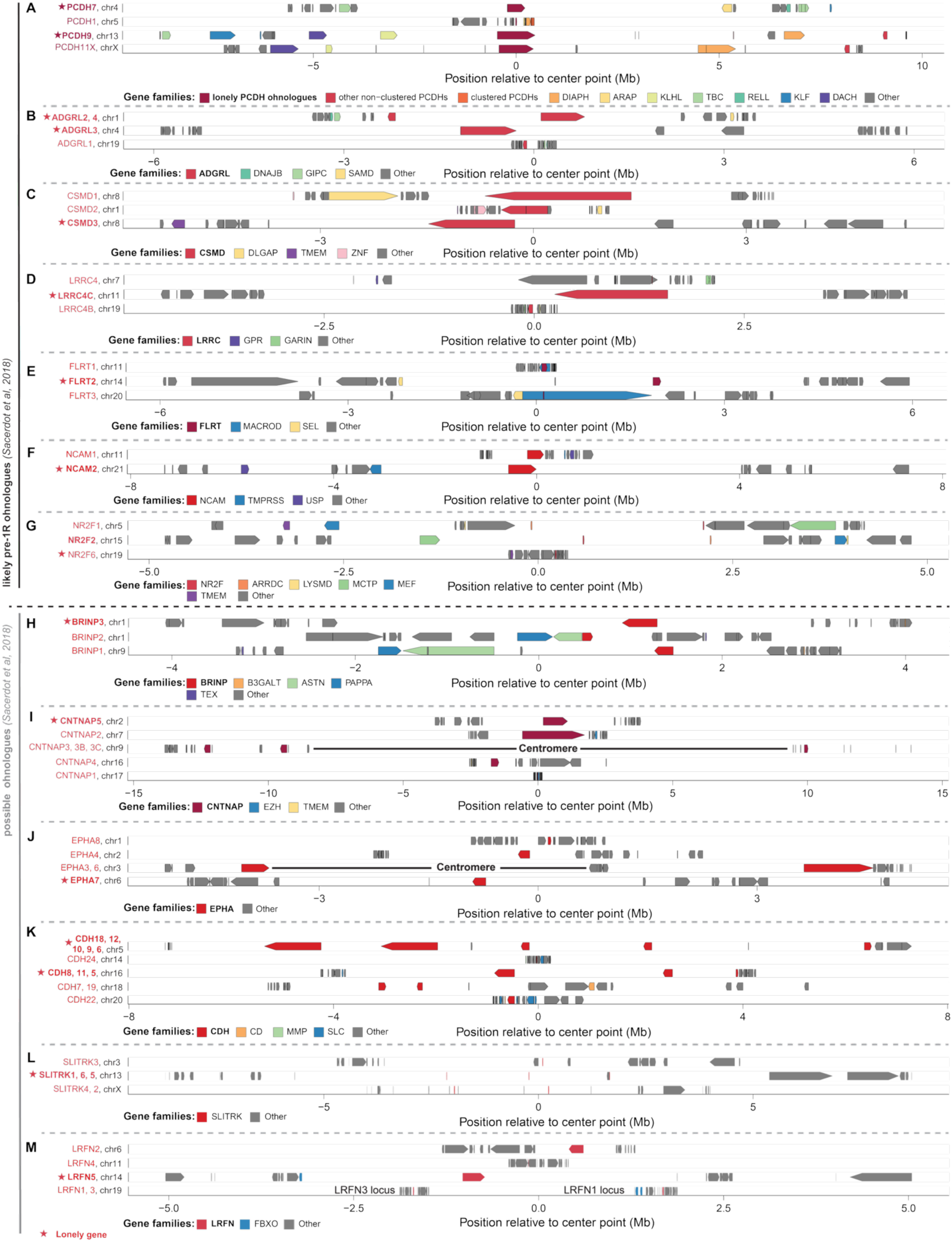
Ohnologs of lonely genes are also often located in or near gene deserts. A-M) Each of these panels display a lonely gene, highlighted in red with a star, and its set of ohnologs in human. Each row displays a locus for one ohnolog relative to the position of the lonely gene. When members of a gene family are present in more than one ohnolog locus, they are colored. The split between A-G and H-M represents the difference in strength of ohnolog calls from Sacerdot et al., 2018 (Sacerdot et al. 2018).

**Figure S4:**
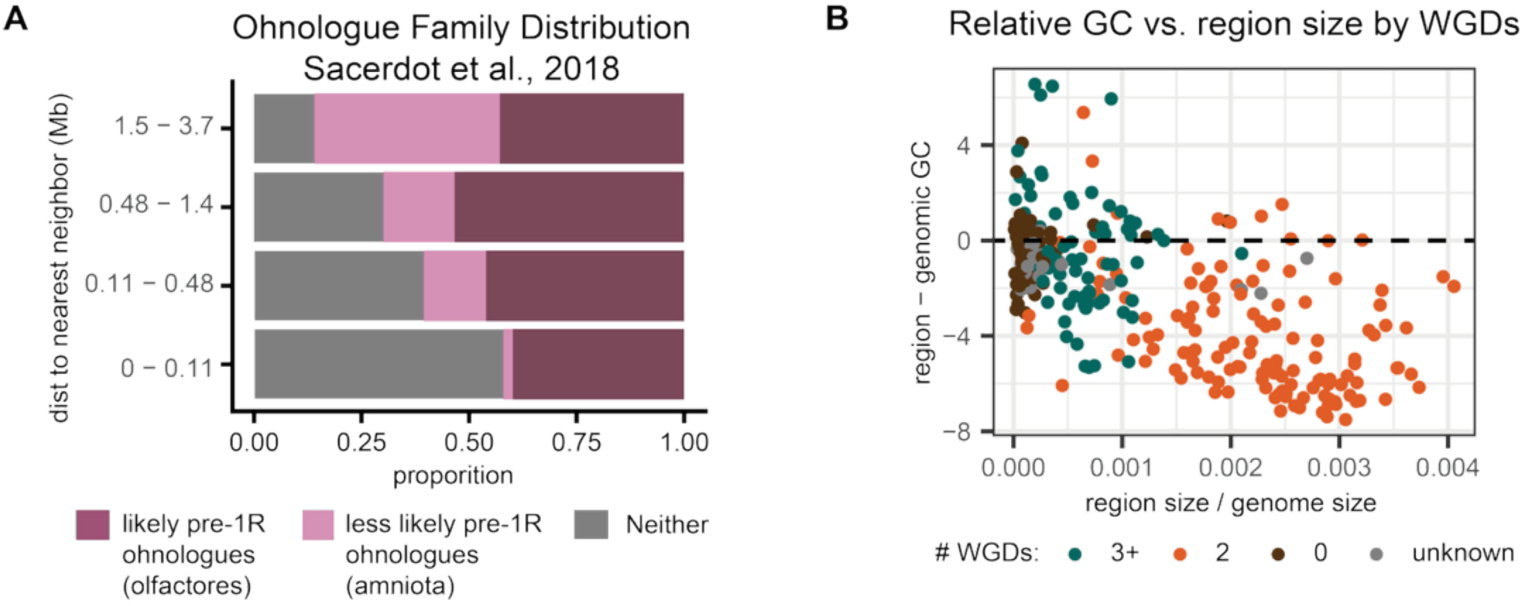
Lonely gene deserts are linked to whole genome duplication events. A) Groups of paralogs predicted to be ohnologs resulting from vertebrate 2R WGDs are displayed according to calculations by Sacerdot *et al*. (Sacerdot et al. 2018). Dark plum represents the proportion of genes in a “distance to nearest” gene cluster that are most likely to have originated as ohnologs. Pink represent Sacerdot’s lower-confidence, candidate ohnologs based on their arrangement in the common ancestor of amniotes. B) Same graph as figure S3F, with dots re-colored to indicate the number of whole genome duplications the source species is thought to have experienced. Relative GC (region minus genomic) versus relative region size (region/genome size) is shown for several lonely gene deserts across metazoans (NCAM2, PCDH7, PCDH9, ADGRL2, ADGRL3, FLRT2). Each point represents a different lonely gene desert, and colors indicate the number of whole genome duplications the source species is thought to have experienced. As in figure S3F, each species can have up to six dots.

Plotting each lonely gene, its predicted set of ohnologs from Sacerdot *et al*., and the closest 10 upstream and downstream genes, we see consistent signatures of microsynteny to support these assignments (Fig 4). For example, the lonely PCDH ohnolog set shares many of the same adjacent gene families across different chromosomes (ie: DIAPH, KLHL, and DACH). Only three genes (GBE1, POM121L12, and SLITRK1) are unlikely to be have ohnologs by Sacerdot’s calculations (Fig. S4A). GBE1 is the only one of its kind, and, as we have noted in a previous section, POM121L12 is a much younger gene, explaining their exclusions (Fig S2B and S3H). SLITRK1, however, resides within a tandem array of SLITRK paralogs, which likely arose through local duplications (in cis) after 2R, rather than through chromosome or segmental duplications. This expanded genomic context probably confounds the ohnolog detection methods, rather than reflecting the true origin of SLITRK1 itself. In fact, the confounding issue of segmental/chromosome versus cis duplication is prominent in several of the lower-confidence ohnolog families (Fig 4I-M).

Large gene deserts (>700 Kb) abut more than one of the predicted ohnologs in most sets (Fig. 4A–C, F–M), yet no set contains a gene desert at every paralogous locus. The most parsimonious explanation is that the desert organization of many lonely genes predates *at least* one round of WGD. The alternative, that each paralogous region independently acquired a similarly massive gene desert, is sufficiently improbable to discount. The remaining 2 lonely ohnolog sets, containing LRRC4C and FLRT2, each have 1 missing ohnolog, making it difficult to determine when these deserts arose. Overall, it is probable that some of the lonely gene deserts arose from whole genome duplication, and it is probable that many of the lonely gene deserts were present before at least one of the two whole genome duplication events at the origin of vertebrates.

### What species have lonely genes, and are they exclusively the genes found in humans?

While the organization we observe in human is therefore ancient, likely preceding at least one round of WGD at the root of vertebrates, we wondered about the possibility of other lonely genes. In other words, are there additional lonely genes in other species that aren’t represented in humans, or is the human repertoire representative of the typical constituents of lonely deserts? To test this, we calculated the “distance to nearest gene” for protein-coding genes in a swath of representative metazoan species. To ensure apples-to-apples comparison across species, we used gene calls available on NCBI for these comparative analyses. These gene calls are less curated than the MANE set we used in human and contain more genes, many of which are not validated. For example, comparing between MANE and NCBI for human, using NCBI we call 44 genes as lonely, including 15 of the 21 we called using our curated set (Supplementary Table 4). The 6 genes only called with our curated list have a nearer neighbor among NCBI gene calls, though they are still quite isolated, and those that are only on NCBI were mostly either 1) unvalidated gene calls excluded from our curated list, or 2) still very isolated, even if not in the most extreme group. To consider a species to have lonely genes, we looked for the following characteristics in the top “distance to nearest neighbor” clusters: 1) all are gene deserts > 250 Kb, 2) at least one lonely gene must be present on an autosome, and 3) the top cluster can’t contain more than 50 genes (ie: to identify true outliers).

Performing this analysis across other metazoan species, we recapitulate the gene-wise analyses represented in Figure 3: proportional to the size of their genomes, most vertebrates harbor lonely genes, and most non-vertebrates do not, excepting cephalopods (Fig 5C, Supplementary Table 5). In species like lancelet (a basal chordate) and tick (Arthropoda: Arachnida), there are still some genes that stand out as more isolated than other genes, but the proportion of the genome these occupy is modest compared to vertebrates or cephalopods.

**Figure 5:**
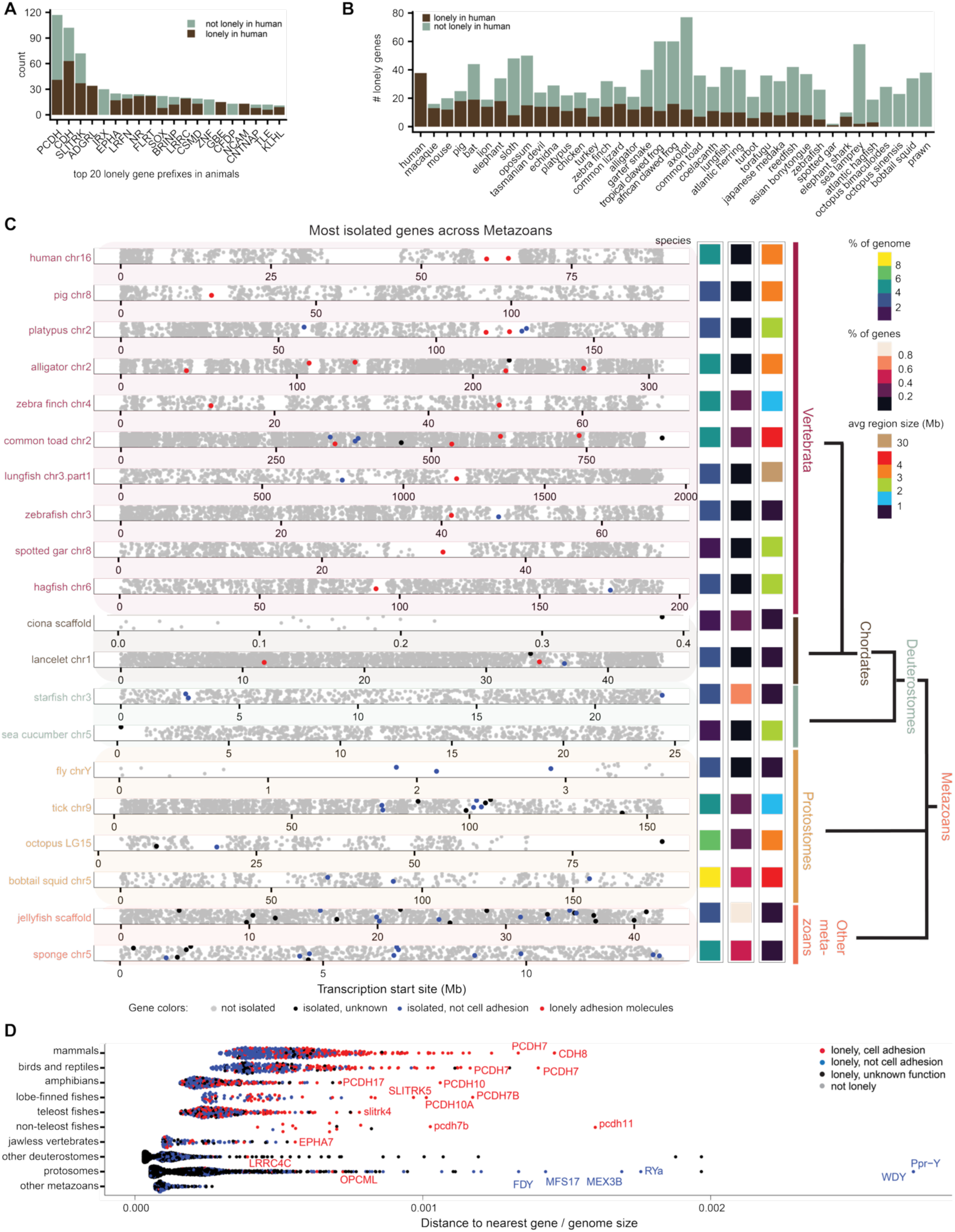
Lonely genes are often cell adhesion molecules across vertebrates. A) Top 20 gene prefixes that are found in lonely genes across animals. Brown shading represents individual orthologs within the protein family that are also found in the human lonely gene set. B) The number of lonely genes is displayed for each species. Colors correspond to the proportion of the set from that species whose human ortholog is lonely. C) Each row displays the chromosome that contains the most isolated gene from that animal. Every gene is represented based on its TSS and colored based on loneliness and function. Lonely genes that are also known cell adhesion molecules are colored red. Note that coelacanth appears less lonely than it is due to the enormity of the chromosome. Chiclets at right display trends for the whole lonely set of genes in that species, as indicated. D) The loneliest genes were collected from each species, as calculated in (A), and then pooled within clades as indicated. Again, colors represent loneliness and cell adhesion.

Interestingly, we do also identify lonely genes in prawn (an arthropod in the crustacean family), despite it not sharing any of the same lonely genes. If we compare the gene spread of the most isolated genes in other species to how isolated they are in humans, we see the most similarity from mammals to human, and the next-most similarity from other vertebrates to human (Fig 5B). In protostomes, there are very few isolated genes that are shared with humans and that shared the same degree of isolation of that protostome species, which may reflect the extensive synteny scrambling that is thought to have occurred in many protostome lineages compared to the bilaterian ancestral gene organization (Schultz et al. 2024). In contrast, vertebrate synteny is thought to retain hallmarks of ancestral bilaterian organization. While we did not find a shared linkage group between the PCDH- and ADGRL-family genes found next to massive gene deserts in cephalopods and vertebrates, we note that the octopus lineage does not share the same degree of synteny scrambling as other protostome clades (Schultz et al. 2024) (Supplementary Table 6-7).

Are the lonely genes that are unique to other species similar to those we observe in humans? Searching by gene prefixes with at least 6 occurrences to select gene types that are occurring in multiple instances across species (ie: more likely to be conserved), we find many of the same gene families occurring over and over again, despite being annotated as paralogs (not orthologs) of those that are lonely in human: cadherins (CDHs), protocadherins (PCDHs), slit and ntrk-like proteins (SLITRKs), bone morphogenic protein/retinoic acid inducible neural-specific proteins (BRINPs), ephrin receptor type A’s (EPHAs), and leucine-rich repeat and fibronectin type domain containing genes (LRFNs) (Fig 5C). We spot-checked the synteny of several of these to ensure they weren’t actually lonely genes being miscalled as paralogs, and we confirmed that they were indeed paralogs rather than orthologs. Of the other gene families, many appear in the second most lonely category in human: Iroquois homeoboxes (IRX), zinc fingers (ZNF), TAFA chemokine like family members (TAFA), and leucine rich repeat transmembrane neuronal proteins (LRRTM) (Fig 5B). Since several of these gene paralogs likely resulted from whole genome duplication, this supports the idea that the non-coding spaces surrounding lonely genes arose before whole genome duplication, and only some of the resulting ohnologs retained the pure desert organization. In others, new genes may have arisen in the desert or infiltrated it, the gene could have been re-located due to large-scale reorganization, or desert sequence may have been lost.

To summarize, if we look at the most isolated genes within vertebrate genomes, we consistently see members of the same gene families occurring on their own in large deserts (Fig 5E). Outside of vertebrates, we do still find some species that have single genes within large gene deserts. These genes can be cell adhesion molecules, but they are less likely to be. This suggests that there is some unique relationship between these gene families and gene deserts in vertebrates, and/or that gene isolation instances observed in invertebrates likely arose separately from those we find in vertebrates.

### Lonely gene deserts exhibit classic features of poorly conserved gene deserts

Given that this genome architecture is ancient within vertebrates, we wondered what could be harbored in these gene deserts that could be permitting, begetting, or maintaining their structure. To ensure subsequent results were representative of lonely gene deserts specifically, rather than of gene deserts or AT-rich zones generally, we created control sets of genes and genomic regions for comparison: 1) centromeres, 2) empty AT-rich gene deserts (i.e. lacking a protein-coding gene in the middle), 3) “almost lonely” gene deserts (ie: genes in smaller gene deserts), 4) AT-rich gene clusters, 5) lonely gene paralogs with some non-coding genic space around them, and 6) GC-rich and almost lonely gene deserts (Fig S6A). Our whole genome calculations for humans were performed across the set of human genomic isochore units, which we computed in previous work (Brovkina et al. 2023).

First, we wondered if lonely gene deserts were unique in their repetitive element composition. To our surprise, their overall repetitive element content was on par with that of the rest of the genome and other control groups, excepting GC-rich deserts (Fig 6A). Like most of our control sets, lonely gene deserts were enriched with LINEs, LTRs, simple repeats, and low complexity repeats, while they were depleted in SINEs compared to genomic background (Fig 6A and Fig S6B). These patterns generally followed AT/GC sequence content. Of these observations, only low complexity and simple repeats were exclusively enriched in lonely gene deserts, though LTRs were specific to the 2 largest gene desert groups – lonely gene deserts and empty, AT-rich gene deserts. Looking into enrichment of specific LTR families, we found ERVL-MaLR’s to be exclusively enriched in lonely gene deserts, while other families were enriched in both lonely gene and classic (i.e., empty) gene deserts (Fig S6C). The high degree of association between these two groups suggests that extremely large, AT-rich gene deserts are more similar to one another whether or not they contain a lonely gene, and they are distinct from smaller gene deserts and other large AT-rich zones.

**Figure 6:**
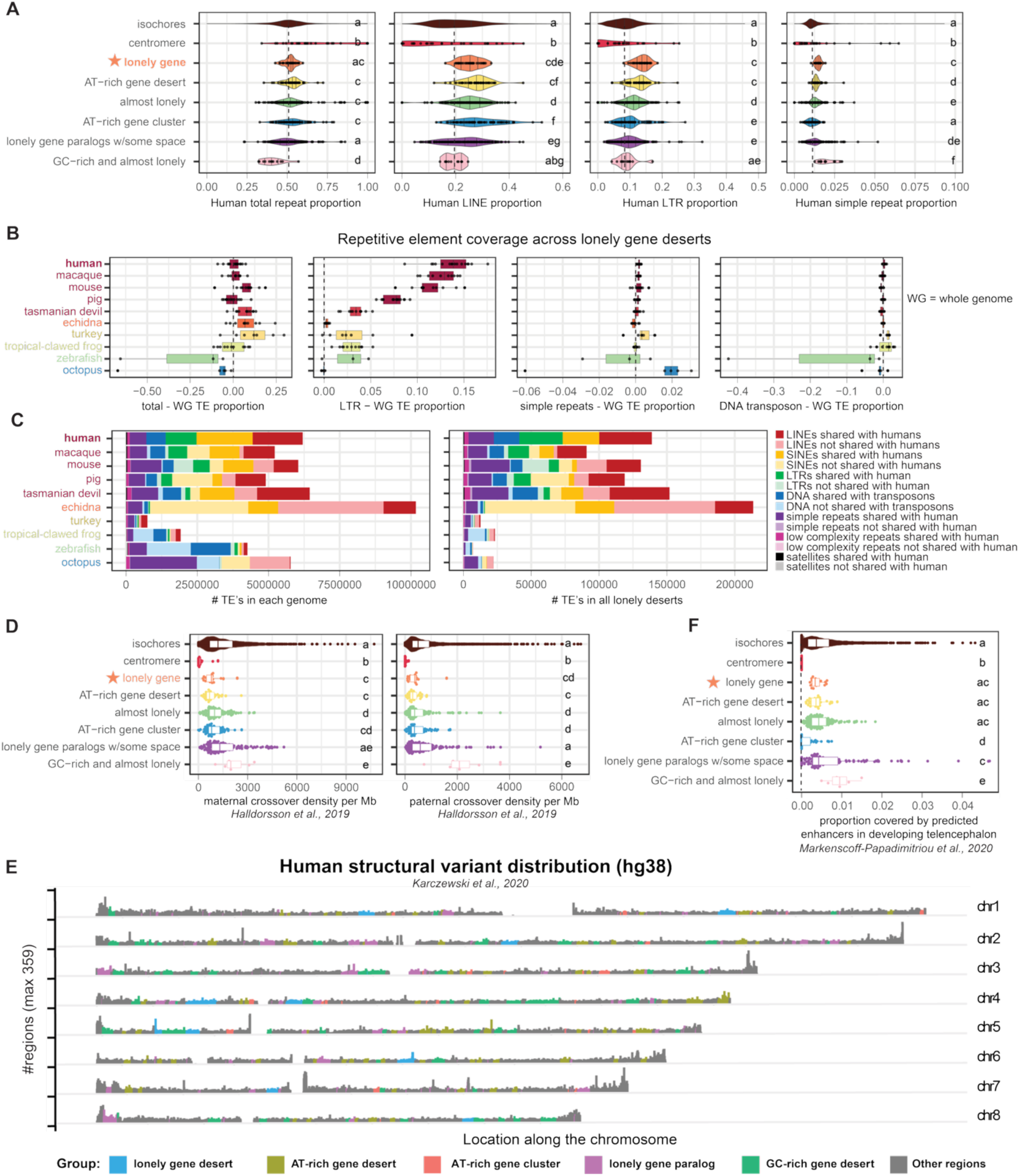
Lonely desert content is variable. A). Human LINE, LTR, simple, and total repeat proportions for lonely regions compared to controls are displayed here. Lonely genes are emphasized in orange with a star. The elements more enriched in the lonely genes are shown here, and other categories of repetitive elements are displayed in Figure S6B and C. B) For each species, lonely gene repetitive element proportion was subtracted from whole genome proportions. Species are colored by clade. The elements more enriched in the lonely genes are shown here, and other categories of repetitive elements are displayed in Figure S6E. C) Bars represent the numbers of each type of element in the whole genome (left) and all shared lonely deserts (right). Darker versions of the same color indicate shared elements, and lighter shades indicate clade-specific elements that are not found in human. D) Maternal and paternal crossover density across lonely genes and controls are displayed here. E) Structural variant distribution across a few representative human chromosomes is displayed here, colored by region. F) Predicted enhancer coverage in the developing telencephalon is displayed across regions. *For all graphs, different letters indicate significant differences as determined by pairwise Wilcoxon test using FDR correction (p < 0.05).

**Figure S6.**
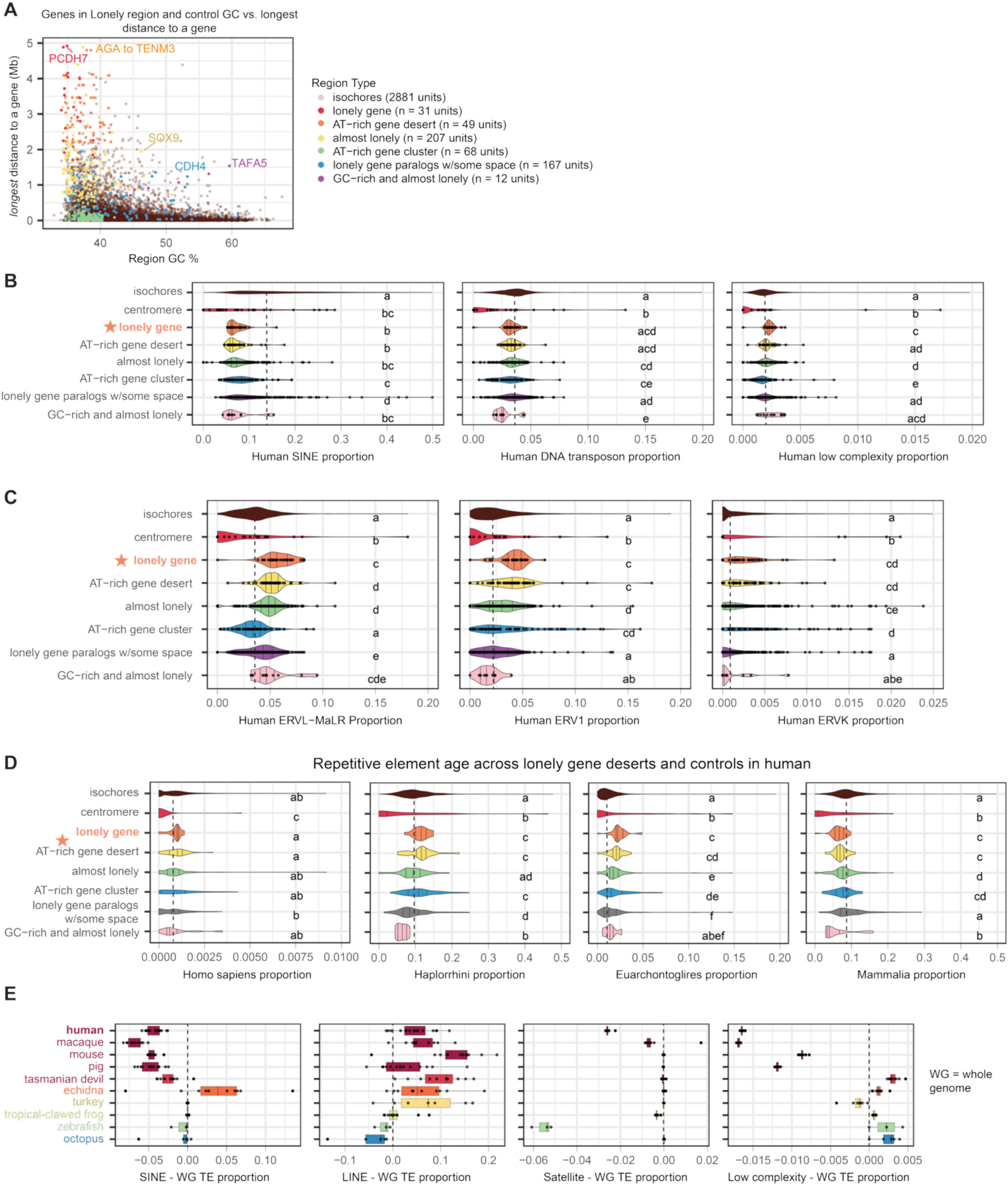
A) Characteristics of lonely gene control regions. For each control set, individual representatives are displayed by distance from the furthest gene and region GC, colored by each region type. The number of genes included in each group are listed in the plot legend. B) Human SINE, DNA, and low complexity repeat proportions for lonely regions compared to controls are displayed here; additional comparisons are shown in Figure 6B. C) Human lonely gene versus control proportions of a few LTR families are displayed here: ERVL-MaLR, ERV1, and ERVK. D) The proportion of each region covered by elements of the following age are displayed: *Homo sapiens, Haplorrhini, Eucarchontoglires,* and *Mammalia*. E) For each species, lonely gene repetitive element proportion was subtracted from whole genome proportions. Species are colored by clade. *For all graphs, different letters indicate significant differences as determined by pairwise Wilcoxon test using FDR correction (p < 0.05).

Despite the structural conservation of lonely gene deserts, we see depletion of older elements and an enrichment of younger elements in these regions, suggesting that they may be experiencing a high degree of TE turnover (Fig S6D). To compare across species and assess if lonely gene deserts contain different types of TEs, we ran RepeatModeler and RepeatMasker on 10 representative bilaterian genomes, and calculated coverage of main repetitive element classes versus whole genome in homologs of human lonely gene regions (Flynn et al. 2020; Smit, Hubley, and Green n.d.). Here, we most consistently find elevated LTR content relative to whole genome, and we also see elevated LINEs, elevated simple repeat, and reduced SINEs in most clades (Fig 6B, Fig S6E). Since this trend was conserved, we wondered if specific elements were consistently present across species. To assess this, we asked how many of the same elements were found in both human and non-mammalian species. As expected, there were progressively fewer shared elements in less related species, with the majority of common elements being simple or low complexity repeat (Fig 6C). In mammals, there was more consistency across other types of element types, but the ratio of new to shared elements appeared around the same relative to whole genome.

If the size and AT content of these regions is consistent over evolutionary time, but specific sequence is in flux, are they experiencing particularly repressed recombination, and as such, are they especially unlikely to exhibit structural variation in human populations? Here, we used crossover data we previously compiled from Halldorsson *et al*. to calculate crossover density in lonely regions (Halldorsson et al. 2019). We find similarly reduced maternal and paternal crossover rates in lonely regions, AT-rich gene deserts, and AT-rich gene clusters other genomic regions (Fig 6D). These data are on par with previous findings that AT-rich zones of mammalian genomes experience reduced recombination, but these findings are not specific to deserts that contain a lonely gene (Brovkina et al. 2023; Duret and Arndt 2008; Duret and Galtier 2009; Waterson et al. 2005).

Compiling structural variants from gnomAD, we see that structural variation in these regions is similar to the genome-wide average (Fig 6E) (Karczewski et al. 2020). Centromeres, pericentromeres, telomeres, and subtelomeres experience the highest rates of structural variants, and the rest of the genome, including lonely regions, experience very few in comparison. Nevertheless, lonely deserts can still experience structural variation.

We describe above that we see loss of desert surrounding lonely genes in genomes that have undergone WGD events, suggesting that a duplication event allows for the relaxation of the constraints holding such large non-coding regions in place. For this reason, we wondered if these regions could be harboring a large number of enhancer elements needed in cis for the regulation of the lonely gene that the genome can’t safely tease apart until there’s a “backup” copy. To address this, since the majority of lonely genes are expressed in the brain, we calculated enhancer density across lonely gene deserts and controls in developing human telencephalon (Fig 6F, (Markenscoff-Papadimitriou et al. 2020). In this tissue, we did not find lonely gene deserts to be significantly enriched for enhancers relative to genomic background or control regions of similar AT content. GC-rich genes in “almost” lonely locales, however, exhibited elevated enhancer density. Several of the genes in these locales are transcription factors (ie: IRX2-IRX4-IRX1 cluster, MSX2, TLE3), so this finding follows previous work illustrating a connection between transcription factor gene deserts and enhancers (Abassah-Oppong et al. 2024; Bagheri-Fam et al. 2006; Huppi et al. 2012; Touceda-Suárez et al. 2020). For these regions, transcription factors are thought to be entangled with nearby deserts that contain their regulatory elements. Our results suggest that this model does *not* apply to the much larger deserts surrounding lonely adhesion genes.

### Lonely genes associate with the nuclear periphery in olfactory epithelium

If the sequence content of the extreme lonely region deserts appears to be ‘normal’ relative to other regions with low gene density and high AT content, then what is the space being used for? To test if it is being used as a structural element, we performed DNA fluorescent *in situ* hybridization (FISH) in adult mouse olfactory epithelium on lonely genes to see where they are located in the nucleus. We chose olfactory epithelium (OE) for this analysis since we have already defined the nuclear architecture of this tissue type (Clowney et al. 2012). To tease apart differences in genomic organization versus expression, we selected probes for pairs of lonely genes and their non-lonely paralogs across a range of expression levels in the OE (Monahan, Horta, and Lomvardas 2019) (Fig 7A and Fig S7).

**Figure 7:**
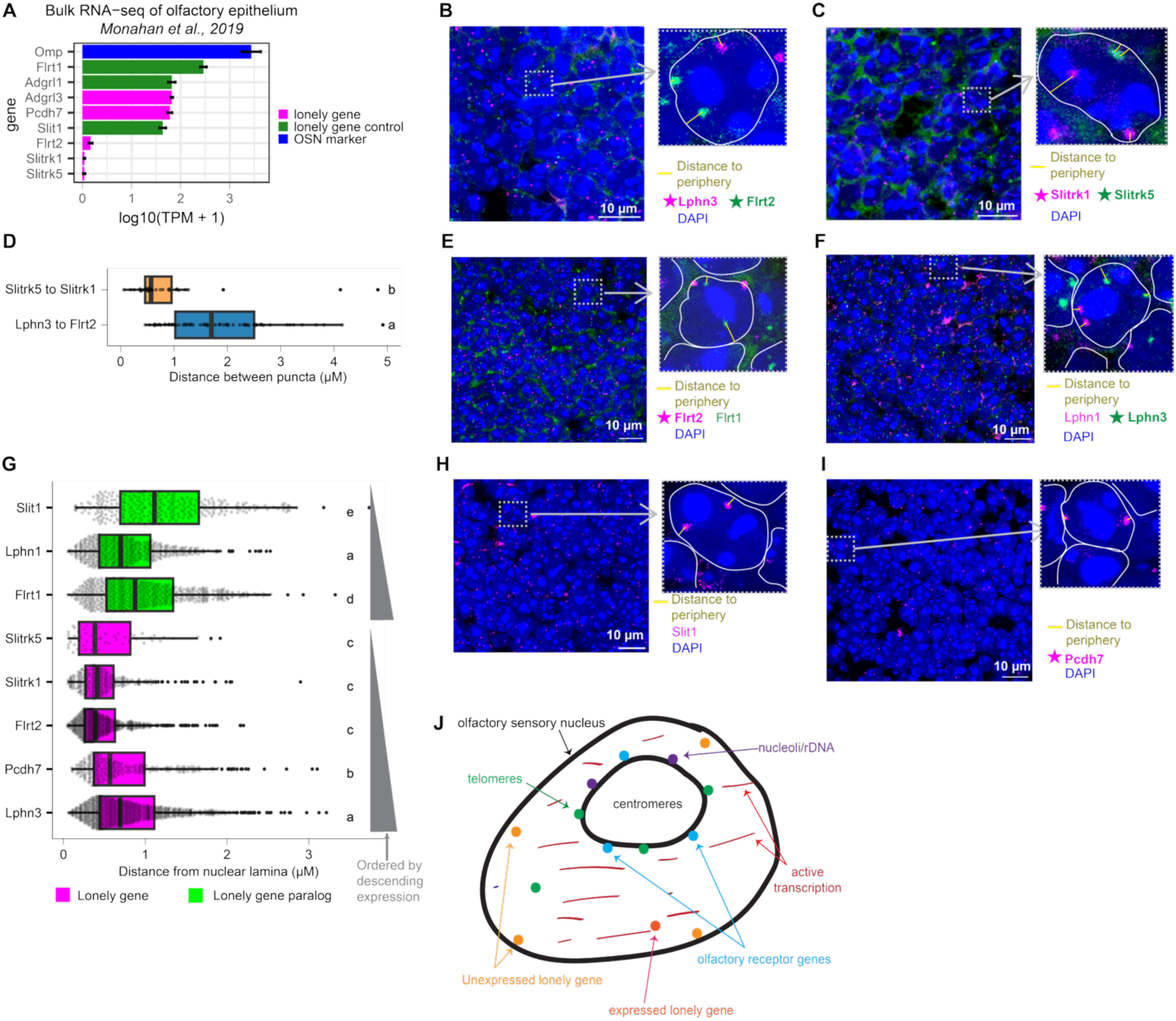
Lonely genes appose the nuclear periphery. A) Bulk RNA-seq of mouse olfactory epithelium from 2 sources is displayed for lonely genes, lonely gene controls, and olfactory sensory neuron marker, Omp (Monahan, Horta, and Lomvardas 2019). B-C, E-F, H-I) Representative DNA FISH images are displayed here for experiments probing 2 lonely genes in one nucleus (B and C) and experiments probing a lonely gene and a control gene E, F, H, and I). The graphical insets indicate how nuclei were identified (circling DAPI signal) and how distance to the nuclear periphery was drawn (ie: from the center of the signal to the nearest nuclear edge). D) Distance between puncta is displayed for each experimental pair. This was measured as the shortest distance from the middle of puncta. Flrt2, Lphn3, and Slitrk5 are lonely; Slit1 is a non-lonely paralogue of Slitrk5. G) Here, we merge all distance to nuclear periphery measurements and order the genes based on expression. Lonely genes are colored magenta and non-lonely paralogous controls are in green. Each group (lonely vs. paralog) is ordered by expression presented in A, as indicated by grey wedges. J) This model of the olfactory sensory neuron nucleus displays locations of several known components of the OSN nucleus (ie: telomeres, olfactory receptor genes, etc.). We show different locales for expressed v. unexpressed lonely genes. *For all panels, different letters indicate significant difference as determined by pairwise Wilcoxon tests using FDR correction (p < 0.05).

Performing DNA FISH experiments on both 1) lonely genes paired with other lonely genes (Fig 7B-C) and 2) lonely genes paired with lonely gene paralogs (Fig 7E-F, H-I), we find that lonely genes tend to be closer to the nuclear periphery than their non-lonely paralogs (Fig 7E-I). We do find lonely gene paralogs to be sometimes associated with the nuclear periphery. Additionally, lonely genes with the highest expression levels in this tissue, Latrophilin-3 (Adgrl3 or Lphn3) and Protocadherin-7 (Pcdh7), stray from the nuclear periphery more than their unexpressed counterparts. Lonely gene paralogs, however, did not display an association between expression levels and proximity to the periphery (Fig 7G).

**Figure S7:**
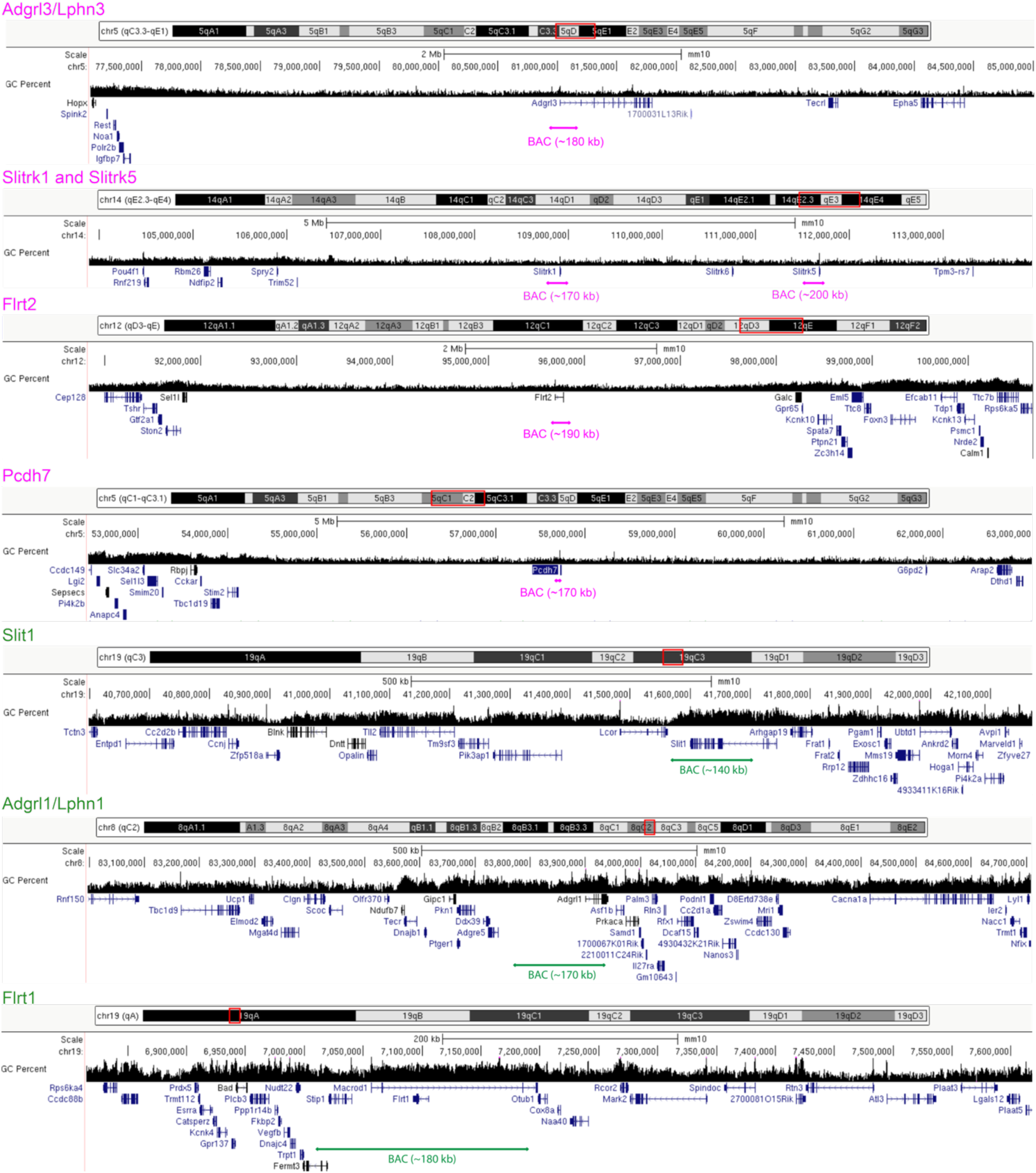
UCSC Genome Browser screenshots showing the BAC probes used for DNA FISH experiments.

## Discussion

The origin and function of gene deserts has long been an area of interest since their surprising discovery in early mammalian genome sequencing work (Gregory 2005; Nobrega et al. 2003; Ovcharenko et al. 2005; Venter et al. 2001). Despite the long history of this field, here we report a novel class: 21 genes in the human genome that are isolated in massive gene deserts. We find that these genes have retained their “lonely” arrangement since early vertebrate history. All but one of these genes is a cell adhesion molecule, associated with ASD, or both. This is distinct from gene desert research to date, which has largely focused on gene deserts neighboring transcription factors (van Steensel and Belmont 2017; Touceda-Suárez et al. 2020). Notably, transcriptional regulators of lonely genes are themselves linked to NDDs. Lonely gene deserts, while structurally conserved in most vertebrates, are lost in a few select subgroups, each of which has their own whole genome duplication that is separate from the 2R WGD events at the origin of vertebrates. Further, DNA FISH in adult mouse olfactory epithelium shows that lonely genes tend to be located closer to the nuclear periphery than their non-lonely paralogs. Activating genes from such a nuclear context may thus require dedicated chromatin remodelers, as was recently shown with MeCP2 in the developing brain (Rullens et al. 2025). It is also possible that the lonely gene arrangement precipitated as a protection from misexpression of the adhesion genes that tend to live in such arrangements, as misexpression of these genes has been shown to be linked to cancer and neurodevelopmental disorders. Below, we discuss the implications of these findings further.

### Are cell adhesion molecules prone to being consumed by gene deserts?

Cell adhesion genes exhibit exquisite cell type specificity in their expression, especially in the brain, which may engender a reliance upon extra sequence space to encode the keys to each cell-type kingdom (Canzio and Maniatis 2019; Krishna-K, Hertel, and Redies 2011; Moreland and Poulain 2022; O’Sullivan et al. 2012; Punovuori, Malaguti, and Lowell 2021). We found that a surprising ¾ of the lonely genes were classed as cell adhesion molecules, a pattern that we observed across vertebrates, despite variation in the exact desert sizes. It is possible, then, that these genes are especially equipped to handle existing in such harsh environments. In the developing brain, for example, cell adhesion molecules are expressed at specific time points in specific cell types during early circuit formation (Flaherty and Maniatis 2020; Moreland and Poulain 2022; Punovuori, Malaguti, and Lowell 2021; Schroeder and de Wit 2018). For a gene of this kind, encasing these switches in heterochromatin could ensure that the proper switch is activated at the proper time, preventing inappropriate or ectopic expression that could result in misdirection of axon guidance cues, and thus, prevent disruption of proper circuit formation. This is supported by numerous studies which illustrate a connection between the lonely genes and both neurodevelopmental disorders and neurodegenerative diseases, though this connection also applies to cell adhesion molecules broadly. Additionally, abnormal expression of cell adhesion molecules, including the lonely genes explicitly, has also been linked to several different types of cancer, including but not limited to glioma and lung cancer (Ando et al. n.d.; Janiszewska, Primi, and Izard 2020; S.-Y. Kim et al. 2011; Mala, Baral, and Somasundaram 2022; Puranik et al. 2024). In this model, adhesion genes transcription is controlled by unusually multidimensional regulatory programs and is particularly stringent; arrangement of these genes in deserts would be a functional adaptation that facilitates exquisite transcriptional control.

### Are the extreme locales of lonely genes a feature or a bug?

This idea does not, however, explain the shear enormity of the lonely gene deserts, nor does it explain the normal enhancer density that we find in these loci. In other words, for a gene to be kept under genomic lock and key does not necessarily mean the gene needs to be smothered by a mountain of intergenic sequence. Olfactory receptors, for example, require specialized transcription factors for their unique expression profile, where only one receptor is expressed in each olfactory sensory neuron, yet these genes live in similarly AT-rich zones, but with comparatively little, extra intergenic space (Brovkina et al. 2023; Chess et al. 1994; Clowney et al. 2011; Monahan et al. 2017; Monahan, Horta, and Lomvardas 2019). Additionally, when olfactory receptor clusters grow too large, they have a tendency to break, and this breakage is hypothesized to be the reason for several chromosomal breaks in mammalian history (J. Kim et al. 2017; Linardopoulou et al. 2005; Mefford et al. 2001; Newman and Trask 2003; Trask et al. 1998; Yue and Haaf 2006). In contrast, we find a high degree of structural conservation in lonely gene desert locales despite variability in sequence content. Adding to the mystery, previous work has shown that deletion of a 1 Mb section of the gene desert surrounding lonely gene ADGRL2 results in few changes to gene expression, and animal survival was unaffected (Nóbrega et al. 2004). So why are these regions so persistent across time?

Perhaps instead, there is a certain percentage of the genome that needs to be comprised of constitutive lamina-associated domains, or cLADs, to maintain the structural integrity of the nucleus, and the lonely gene deserts of mammals supply a large percentage of them. We know from previous work that if the ability to form LADs is depleted from developing neurons by knocking out lamin proteins, the structural integrity of the nucleus is diminished at best, and the animal does not survive long after birth at worst (Alabi et al. 2025; Coffinier et al. 2010, 2011; Shin et al. 2025). In other words, it is possible that the deserts surrounding lonely genes are functional but not especially adaptive for the genes therein; instead, the entrapment of these genes in these conditions was just a side effect, a runaway train around which transcriptional machinery was able to adapt through evolution of chromatin modifiers like POGZ and MeCP2. The primary function of these regions, in this case, is structural, and the deleterious effects we see associated with lonely genes could be a consequence of their entrapment in such harsh conditions.

If this structural case is true, it is possible that these regions are remnants of ancient centromeres. In Fig 1D, we find that lonely gene deserts look remarkably similar to centromeres: they are massive and there is around one per chromosome. Functionally, we also know that the heterochromatin adjacent to centromeres is typically included in LADs, which lonely gene deserts seem to share (van Steensel and Belmont 2017). Furthermore, our data are consistent with an origin of lonely gene organization occurring some time before the 2R vertebrate genome duplication events, events that double centromeres, as well as gene content. If these ancient duplication events resulted in any chromosomes that then fused together, it would generate 2 centromeres in the chromosomes that underwent fusion. Work from Sacerdot *et al* suggest that 7 chromosomal fusion events likely took place after the first vertebrate WGD, and another 4 fusions occurred after the jawless vertebrates branched off. Adding to this possibility, we found that 2 lonely gene ohnologs, CNTNAP2 and EPHA3/EPHA6 were located directly adjacent to a centromere (Fig 4I and K). If these regions are, in fact, old centromeres, enough evolutionary time has passed that we would no longer see repetitive signatures of those centromeres anymore, but perhaps a lingering remnant of their structural role has remained.

### Lonely genes and peripheral nuclear localization

In our DNA FISH experiments, we find that lonely genes tend to localize near the nuclear periphery, particularly when they are not expressed. In olfactory sensory neurons, where nuclear architecture is inverted relative to most other cell types, this peripheral positioning is especially striking (Clowney et al. 2012). These observations suggest that lonely genes frequently reside within lamina-associated or lamina-proximal chromatin environments. Constitutive lamina-associated domains (cLADs) are typically gene-poor and enriched in AT-rich sequence, features that are characteristic of the deserts surrounding lonely genes (Meuleman et al. 2013; van Steensel and Belmont 2017). The large, heterochromatin-prone environments in which these genes reside may therefore impose structural or regulatory constraints on their expression. Several lonely genes have been reported to be sensitive to disruption of chromatin regulators implicated in neurodevelopmental disorders, including POGZ and MeCP2 (Boxer et al. 2020; Li et al. 2020; Markenscoff-Papadimitriou et al. 2021). While the present study does not directly test the mechanisms by which these factors influence lonely gene expression, the enrichment of regulatory sensitivity among genes embedded in massive deserts raises the possibility that genomic isolation creates a form of regulatory vulnerability.

We previously characterized the nuclear organization of ORs (which are AT-rich, paralogous arrays) and many kinds of repetitive elements in the olfactory epithelium (Brovkina et al. 2023; Clowney et al. 2012). While heterochromatinized sequences associate with the nuclear lamina in many cell types, we found that OSNs have an “inverted” nuclear organization, in which chromocenters and other heterochromatinized sequences occupy a “fried egg yolk” in the center of the nucleus. This organization occurs due to lack of lamin B receptor in this cell type. The peripheral location of the lonely genes in this cell type is thus quite striking in contrast to everything we looked at previously. Together, we present a testable model of the olfactory sensory nucleus in which unexpressed lonely genes are tethered to the nuclear lamina and expressed lonely genes are pulled off the lamina when transcribed (Fig 7J). Remarkably, some of the genes we describe as “lonely” have also been found to be associated with the nuclear lamina in cultured cells (Meuleman et al. 2013). More recently, MeCP2 has been hypothesized to allow expression of lamina-associated genes in the developing brain (Rullens et al. 2025). Together, we propose that lonely neurodevelopmental genes are tethered, within their deserts, to the nuclear periphery. Allowing expression from these domains requires the action of specific regulatory proteins, leading to a vulnerability in brain development.

## Methods

### Distance to nearest gene and furthest distance to gene calculations

To identify lonely genes in the human genome, we collapsed the list of protein-coding genes in hg38 down to one isoform per gene. When choosing an isoform, we selected the isoform provided by MANE Select (Matched Annotation from NCBI and EMBL-EBI) first; these coordinates were identified by choosing representative transcripts for each gene by matching exonic sequences in both NCBI and RefSeq (Morales et al. 2022). This set misses about 3% of the genome, so to complete the rest of these sequences, we grabbed each gene’s longest isoform listed in the RefSeq set of protein-coding genes. With this set of ∼20,000 human protein-coding genes, we calculated the distance to nearest gene as the smallest choice between the following:

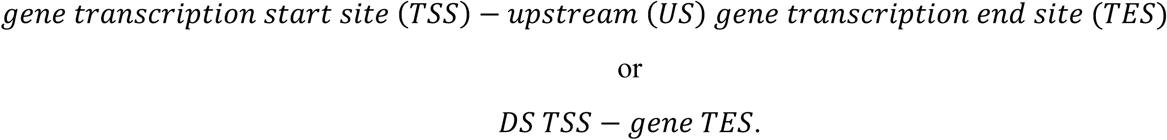

We then performed k-means clustering on these values. Based on an elbow plot (Within-Cluster Sum of Squares (WCSS) vs. k-values), 3 to 5 clusters would have been reasonable. We looked at the top cluster from each of these 3 choices (ie: the most ‘isolated’ genes), and the top genes were the same regardless of the choice; only the bottom cluster groupings shifted. Since the choice didn’t matter for our genes of interest, we selected the middle option, 4 clusters.

We use this same set of protein-coding genes to calculate the furthest distance to a gene, instead selecting the largest choice between the following:

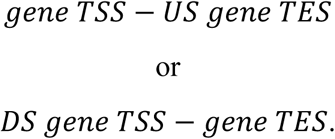

We performed the same k-means clustering technique, and we again selected 4 clusters. For cross-metazoan distance calculations, we collected GFF files from RefSeq for a slew of representative metazoan species, and we filtered these annotations to just protein-coding genes (Supplementary Table 8).

### Cell adhesion annotation

Since there are only 21 lonely genes, we manually searched each of these genes in the literature for identified cell adhesion function. We also cross-referenced UniProt’s list of cell adhesion molecules (same as GO:0007155) with a manual literature search (The UniProt Consortium 2025). Because only 1 of 21 genes was incorrectly not called as a cell adhesion molecule from the lonely set, we simply used AmiGO (cell-cell adhesion, GO:0007155) to call the remaining human genes (Carbon et al. 2009). GO analyses were performed in R using the enrichGO function from clusterProfiler package, with a p-value cutoff of 0.05 and a q-value cutoff of 0.05 using the Benjamini-Hochberg correction (Wu et al. 2021; Xu et al. 2024; Yu 2024; Yu et al. 2012).

### Bulk RNA-seq analysis

Re-bulk RNA-sequencing feature counts were pulled for all analyses, except POGZ, which we generated from fastq files (Markenscoff-Papadimitriou et al. 2021; Monahan, Horta, and Lomvardas 2019; Tenreiro et al. 2025; Wijayatunge et al. 2018). Fastq files were processed using standard workflow and aligned to the mouse genome (mm9) using HiSat2 (HISAT: a fast spliced aligner with low memory requirements | Nature Methods n.d.). Feature counts were collected with featureCounts, then they were processed in DESeq2 (v. 1.44.0), marking significance p < 0.05 and log2FC > 1 (Liao, Smyth, and Shi 2014; Love, Huber, and Anders 2014).

### Manual synteny tracking

For species whose lonely genes we were unable to track in Genomicus, we manually tracked their synteny. To do this, we pulled RefSeq GFF annotation files for a list of representative animal genomes, and we filtered annotations to just protein-coding genes. We then searched for human lonely gene names in each. For any missing gene names, we searched gene descriptions and were able to find lonely gene orthologs for each species without needing to run a separate BLAST. For graphing, we then pulled 10 genes on either side, and we colored each gene that occurred in that locus at least 4 times across species. If a species lost synteny on both sides, we excluded it from analysis, but it was rare that this happened; typically, we either found gene and synteny, or we couldn’t find the gene at all. If there was a partial loss of synteny at a locus, we searched from the missing flank elsewhere in the genome to see if the space had moved with the flank, but we did not find a case of this happening with any of the genes we analyzed.

### Linkage group identification and reciprocal BLAST

To identify linkage groups, we used Oxford Dotplot (odp) from the Simakov Lab (Schultz et al. 2023). We first identified linkage groups between a few select species with known lonely gene deserts and a range of evolutionary distance: human, mouse, and spotted gar. Finding lonely genes in these linkage groups, we performed odp with human, mouse, spotted gar, and East Asian common octopus, both with and without tick. We searched the reciprocal BLAST hits files to identify the presence of an ortholog call for a human lonely PCDH and ADGRL.

To identify orthologs to human genes across metazoans, we used odp, modifying the script to only perform a reciprocal BLAST of each proteome against the human (hg38) proteome, rather than of each proteome against each other.

### Repetitive elements

Here, we ran RepeatMasker and RepeatModeler on a set of representative vertebrates from NCBI (Table 2) (Goldfarb et al. 2025). We calculated coverage of classes, families, and subfamilies separately, using the valr package in R (Riemondy et al. 2017). For the human-specific analyses, control regions are as follows:

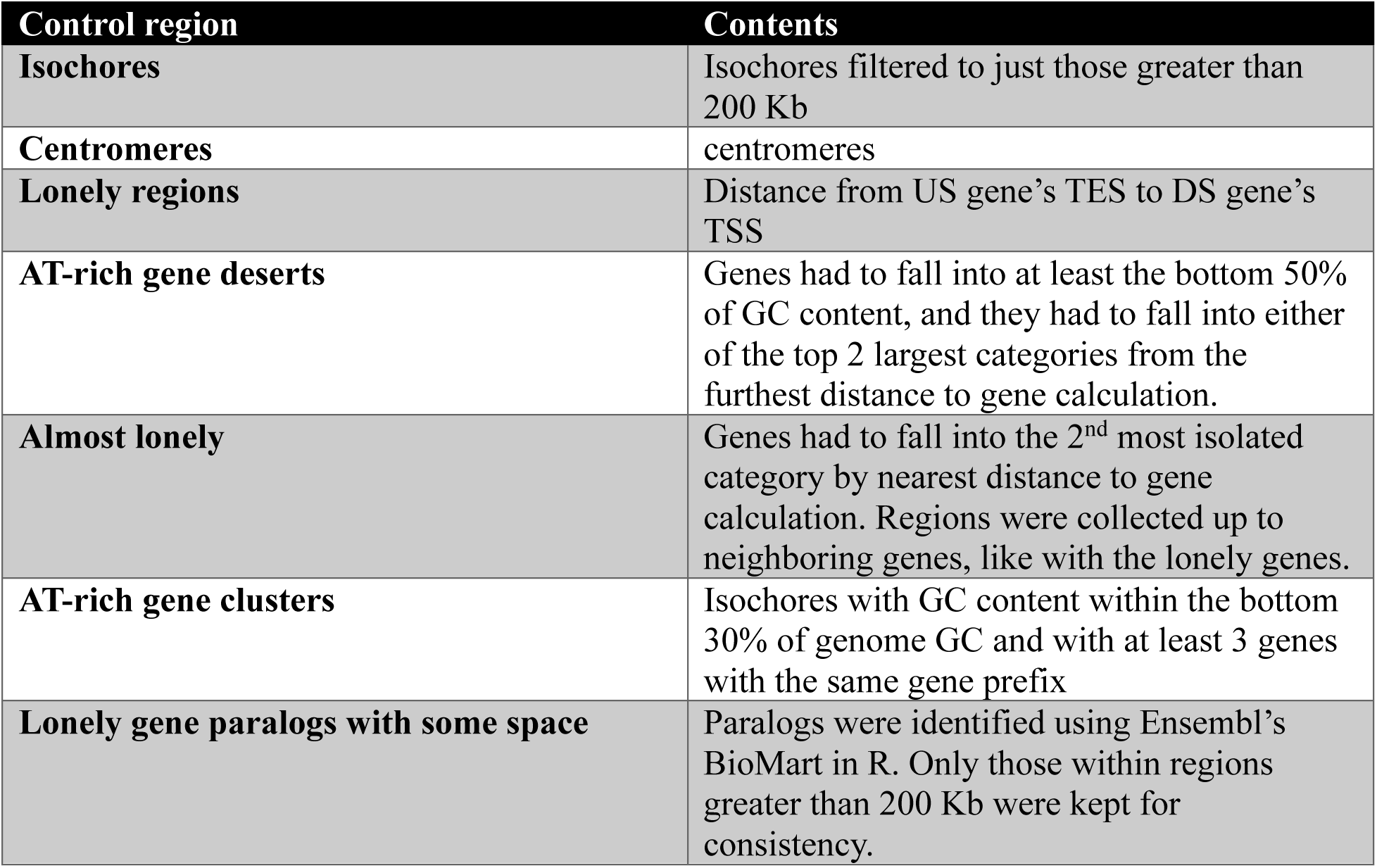
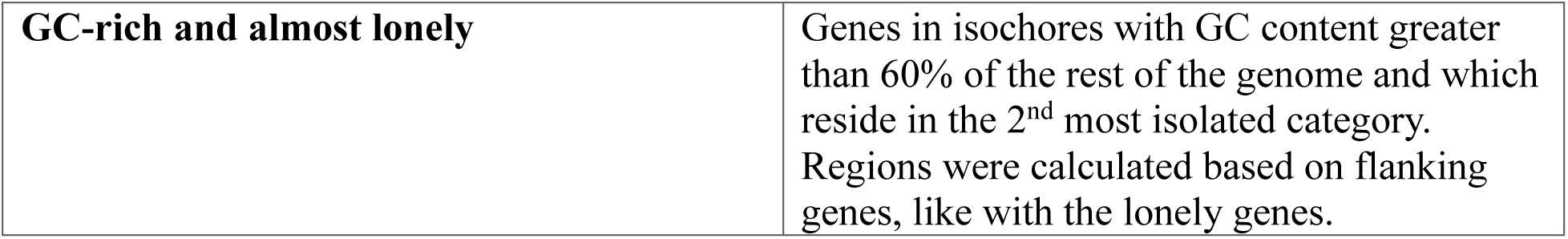

All regions excluded gaps and blacklisted regions, and regions derived from isochores are pulled from Brovkina and Chapman et al (Brovkina et al. 2023). For lonely regions from other species, we pulled genes which were identified as lonely in both human and that species, then grabbed the region between the genes paralogous to the flanks identified in human. In all species, there weren’t genes located between these paralogous genes, other than the lonely gene. TE ages were extrapolated from Dfam (Hubley et al. 2016).

### Recombination and Structural variants

Crossover densities were calculated from data generated by deCODE, in which whole genome sequencing (WGS) was performed on trios, allowing for refinement of 247,942 crossovers in 9423 paternal meiosis and 514,039 crossovers in 11,750 maternal meioses (Halldorsson et al. 2019). Same regions and controls were used here as in repetitive element annotation and coverage calculations. Structural variants were pulled from gnomAD v4.1, a conglomeration of 445,857 SVs from 10,738 unrelated individuals(Karczewski et al. 2020). Here, we filtered to just those variants with strong support to be called (aka: filter = PASS). We included all remaining variants in our final figure, but we got similar results when we reduced SVs to those coming from individuals without neurological disease (non-neuro) and to those just coming from controls.

### DNA FISH and image analysis

DNA FISH was performed on adult mouse olfactory epithelium, as described by Clowney et al, with the following exceptions (Clowney et al. 2012). Here, we collected 8 µM sections, and sections were denatured at 95°C for 8 minutes. Tissue was collected from mice of the 129S1/SvImJ strain, ages 21-28 days. Images were analyzed in ImageJ (Rueden et al. 2017). When measuring distance to nuclear periphery, we turned off the color channel containing the probe, identified the edges of the DAPI signal for a cell, turned the probe’s color channel back on, then measured the shortest distance from the center of a probe signal to the edge of the DAPI signal. For distance between probe signals, we measured the distance between the centers of two different probes within the same cell, favoring the pairings which allowed for the shortest distances.

